# Substrate binding and turnover modulate the affinity landscape of dihydrofolate reductase to increase its catalytic efficiency

**DOI:** 10.1101/2020.04.14.040733

**Authors:** Nicole Stéphanie Galenkamp, Giovanni Maglia

## Abstract

It is generally accepted that enzymes structures evolved to stabilize the transition-state of a catalyzed reaction. Here, observing single molecules with a multi-turnover resolution, we provide experimental evidence for a more sophisticated narrative. We found that the binding of the NADPH cofactor to DHFR induces a first allosteric change that increases the affinity of the enzyme for the substrate. Then the enthalpy generated by the chemical step provides a power stroke that switches the enzyme to the product-bound conformations and promotes the release of the oxidized cofactor NADP+. The subsequent binding of NADPH to the vacated site provides the free energy for the recovery stroke, which induces the allosteric release of the product and resets the initial configuration. Intriguingly, the cycle is not perfect. Occasionally, DHFR undergoes second-long catalytic pauses, most likely reflecting the occupancy of an off-path conformation induced by excess energy liberated by the chemical step. This catalytic remodeling of the affinity landscape of DHFR is likely to have evolved to improve the efficiency of the reaction to cope with the high concentration of NADP+ in *E. coli.* And might be a general feature for complex enzymatic reaction where the binding and release of the products must be tightly controlled.

## Introduction

The mesmerizing power of enzymes to catalyze chemical reaction has fascinated scientist for over a century. The first crystal structure of an enzyme, lysozyme^1,2^, confirmed that enzymes fold into a three-dimensional structure that stabilizes the transition state of the catalyzed reaction. Fifty years on, however, scientists are not yet able to rationally design enzymes. The failure of catalytic antibodies to catalyze reactions with enzyme-like efficiency suggested that transition-state stabilization is only part of the picture. In the meanwhile, other characteristics of enzymes emerged. The first is that enzymes are not rigid structures, and their flexibility and dynamics is an important factor to efficiently catalyze reactions.^3–6^ Another is that enzymes might adopt more than one thermally stable conformation^7–10^, which might be linked to catalysis.^11–13^ And as last another intriguing characteristic of enzymes is that they appear to have large structures stabilized by a network of hundreds of weak interactions, and the disruption of just a few such interactions (sometimes one) can have profound effects on the catalytic efficiency of the enzyme.^14–16^

Dihydrofolate reductase (DHFR) has been used extensively as a model enzyme for investigating the link between structure, dynamics and catalysis.^17–21^ The enzyme catalyses the reduction of 7,8-dihydrofolate (DHF) to 5,6,7,8-tetrahydrofolate (THF) using nicotinamide adenine dinucleotide phosphate (NADPH) as cofactor (**Fig. 1A**). Five intermediates have been identified in the steady-state catalytic cycle^22^ and characterized by X-ray crystallography^23–27^ and NMR^18,20,21,28,29^: E:NADPH, E:NADPH:DHF, E:NADP^+^:THF, E:THF, and E:NADPH:THF (**Fig. 1B**). Large conformational changes are observed in the Met20 loop (residues 9 to 24) between the Michaelis complex E:NADPH:DHF [modeled by E:NADP^+^:folate in structural studies] and the product complex, [modeled by E:NADP^+^:THF].^23^ The Met20 loop adopts two main conformations: the “closed” conformation, in which the loop packs tightly against the nicotinamide ring of the cofactor, and the “occluded” conformation, in which the loop sterically blocks the nicotinamide-binding pocket. The holoenzyme (E:NADPH) and the model Michaelis complex (sampled by E:NADP^+^:folate) are in the closed conformation, whereas the product complexes adopt the occluded conformation (**Fig. 1B**).^18,21,23^

**Figure 1.**
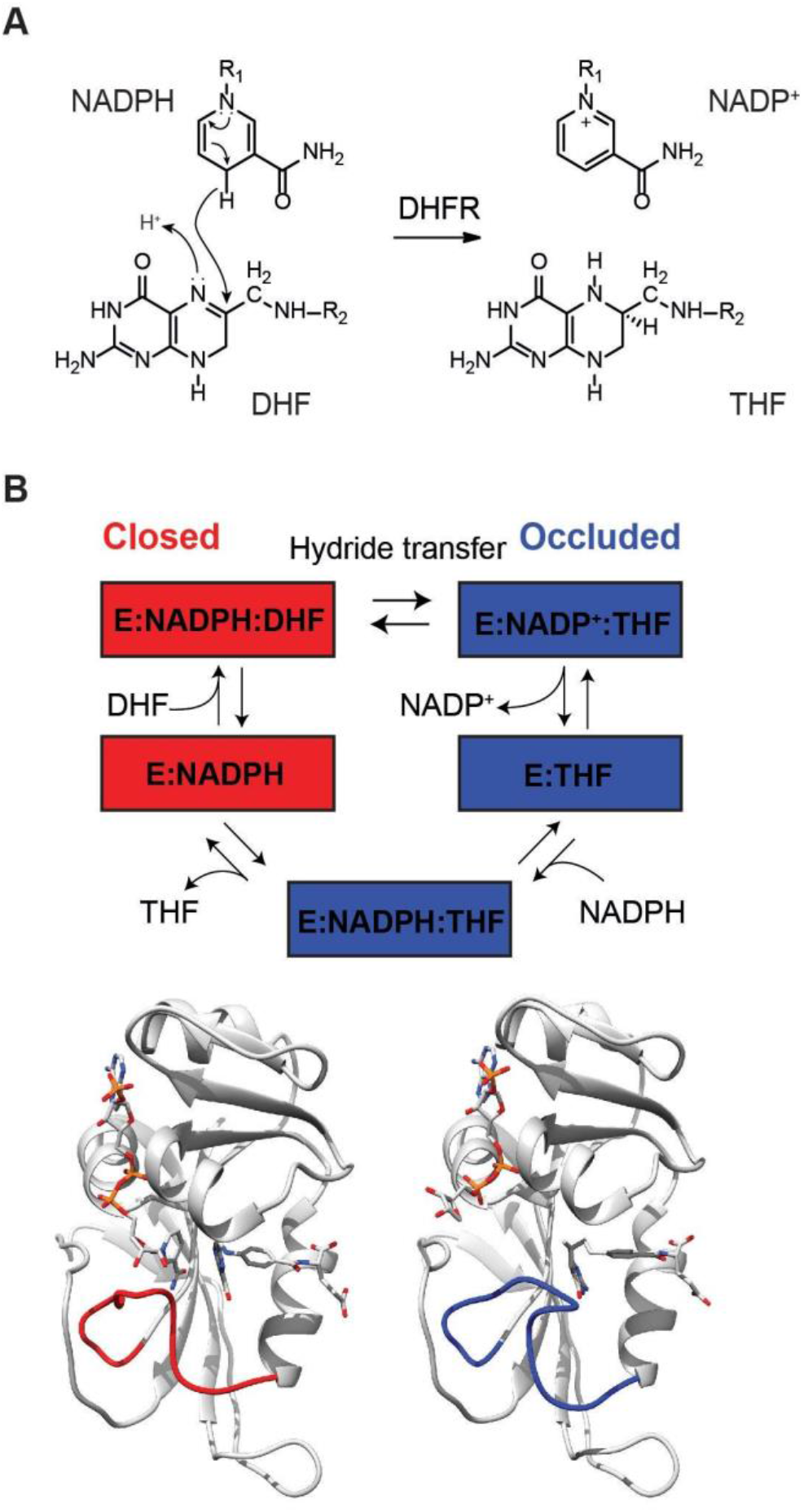
DHFR reaction. (**A**) DHFR catalysed reaction. (**B**) DHFR catalytic cycle as identified by X-ray crystallography and NMR spectroscopy for the Met20 loop conformation. The cycle starts with DHFR in complex with NADPH (E:NADPH, PDB 1RX1), then dihydrofolate binds (E:NADPH:DHF, PDB 1RX2). In both of these structures, DHFR adopts the closed conformation, where the Met20 loop (red loop) is packed against the nicotinamide ring of the cofactor. After the reaction, the enzyme switches to the occluded conformation whereby the Met20 loop sterically hinders the nicotinamide-ribose binding pocket. Release of NADP^+^ (PDB 1RC4) and rebinding of NADPH (PDB 1RX6) precede the release of the THF (PDB 1RX5). Below are structures of the closed ternary complex E:NADP^+^:FOL: (left, PDB 1RX2) and occluded ternary complex E:NADPH:5,10-dideazaTHF (ddTHF)(Right, PDB 1RX6) illustrating the conformational changes that occur upon hydride transfer the M20 loop shown in red and blue, respectively.

In this work, we sample the catalytic reaction of DHFR by ionic current recordings. To trap DHFR inside the nanopore, a cysteine-free DHFR was extended with a positivity charged C-terminal polypeptide tag and two negatively charged residues were introduced to the surface (named here as DHFR_tag_).^30,31^

The long observation time, which allowed recording for the first-time tens to hundreds of turnovers from a single enzyme, unraveled an unexpected mechanism in DHFR catalysis. We previous showed that DHFR_tag_ exists in at least four ground-state conformations or conformers that have different affinity for various ligands^31^. Here we show that the free energy of the chemical step and that of substrate binding is used by the enzyme to promote conformational exchange and modulate the affinity of the enzyme for the reactants and products. We further observe that the catalytic step occasionally induces the formation of an inactive off path conformer in DHFR. This suggests that the soft interactions in folded enzymes might be important not only to provide the ideal chemical environment for the reaction to occur, but also to dissipate the excessive energy generated during the catalysis.

## Results

### Binding of substrate ligands to DHFR_tag_

In nanopore recordings, an external applied potential is used to drive proteins inside a single protein nanopore, which is observed by the step-wise reduction of the open pore current (*I*_O_) to a blocked pore current value (*I*_B_) (**Fig. 2A-C**). Typically, fractional residual current percent (*I*_res%_ = I_B_ / *I_O_* x 100) are reported to account for pore-to-pore variations. Modulations to the ionic current are then used to characterize molecules binding to the trapped enzyme. Importantly, previous work with DHFR_tag_ or other proteins showed that the confinement inside the nanopore, and the effect of the electric field does not affect the thermodynamic stability of the protein within the nanopore.^30,32–37^

**Figure 2.**
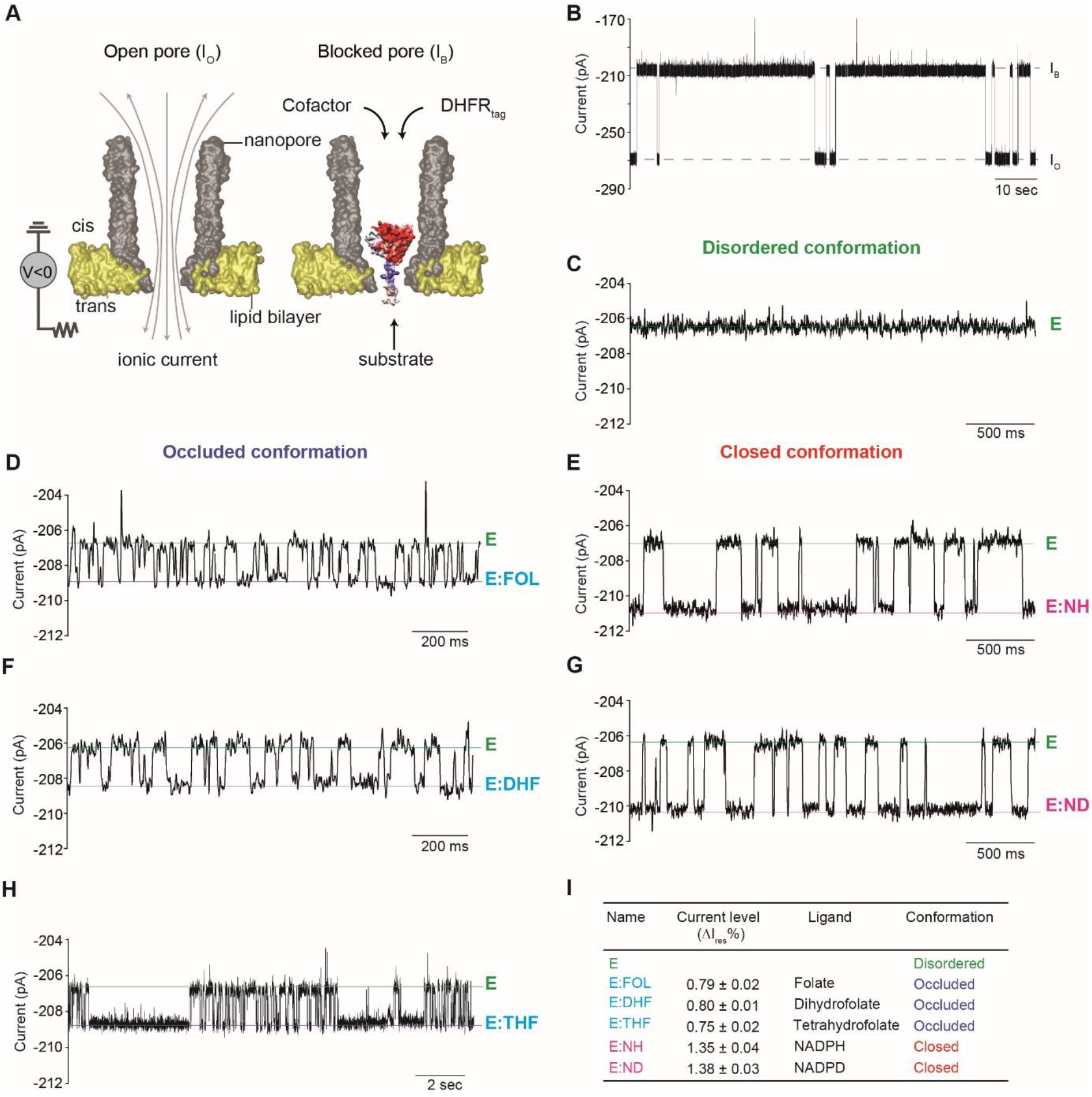
Binding of ligands to DHFR_tag_ inside the ClyA nanopore. **(A)** Left: Surface representation of a ClyA-AS nanopore (grey) embedded in a planer lipid bilayer (yellow). Right: Surface representation of DHFR_tag_, which is colored according to the vacuum electrostatics (Pymol), inside ClyA-AS^30^. **(B)** Typical ionic current blockades provoked by the capture of a single DHFR_tag_ molecule (50 nM, *cis)* by ClyA-AS at −80 mV. **(C-J)** Expansions of typical blockades of DHFR_tag_ in the absence of ligands **(C)** and in the presence of 33.6 μM folate *(trans,* D) 26.6 μM NADPH (*cis*, **E**), 5 μM dihydrofolate *(trans,* **F**), 22.5 μM NADPD *(cis,* G), 12.6 μM tetrahydrofolate *(trans,* **H**). **(I)** Table showing the connection between the observed current levels and the different structure of DHFR as obtained by X-ray crystallography^23^. All current traces were collected in 250 mM KCl, 15 mM Tris-HCl pH 7.8 at 25 °C, by applying a Bessel-low pass filter with a 2 kHz cut-off and sampled at 10 kHz. All traces except the recordings in B were filtered digitally with a Gaussian low-pass filter with a 100 Hz cut-off.

In this work, the different conformations of DHFR were probed by adding the slow-reacting folate (FOL), the substrate dihydrofolate (DHF) and the product tetrahydrofolate (THF) to the *trans* side of the nanopore, and the cofactor NADPH (or its deuterated (NADPD) or reduced (NADP^+^) form) to the *cis* side of the nanopore. NADPH and NADPD bind to the closed configuration of the enzyme^23^, while FOL, DHF and THF bind to the occluded configuration of the DHFR^23,38^ (**Fig. 1**). The binding of ligands induced discrete current enhancement from the basal *I*_res%_ blockades of DHFR_tag_ and are expressed as Δ*I*_res%_ (**Table 1, Fig. 2D-I**). The binding of substrates to the occluded configuration induced lower current enhancement compared to the binding of the cofactor to the closed configuration. From the blockades, the association and dissociation rate constants of all ligands can be measured (**Table 1**). Interestingly, tetrahydrofolate showed two dissociation rates (1.39 ± 0.2 s^-1^ and 27.5 ± 2.0 s^-1^, **Table 1, Fig. 2H**), possibly corresponding to the binding of THF to either the closed and occluded conformation of DHFR. The binding of NADP^+^ to DHFR_tag_ was not observed, because of its fast dissociation constant (*k*_off_ = 300 s^-1^).^22^

**Table 1:** Current blockades and kinetic constants of DHFR_tag_ upon addition of NADPH, NADPD *(cis* compartment), folate, dihydrofolate and/or tetrahydrofolate *(trans* compartment). Errors are given as standard deviation between independent pore (N = 3). All current traces were collected in 250 mM KCl, 15 mM Tris-HCl pH 7.8 at room temperature (25 °C) at −80 mV. (L) and (H) indicate the low- and high-affinity off rates of THF.

| | $I_{\text{res-bound}}$ (%) | $I_{\text{res-apo}}$ (%) | $\Delta I_{\text{res}}$ (%) | $\Delta I_{\text{res}}$ (pA) | $k_{\text{on}}$ ($\mu\text{M}^{-1} \text{s}^{-1}$) | $k_{\text{off}}$ ($\text{s}^{-1}$) | $K_{\text{D}}^{\text{app}}$ ($\mu\text{M}$) |
| --- | --- | --- | --- | --- | --- | --- | --- |
| **Folate** | 75.6 ± 0.5 | 75.8 ± 0.5 | 0.79 ± 0.02 | 2.1 ± 0.1 | 1.2 ± 0.3 | 63.4 ± 3.2 | 53 ± 14 |
| **DHF** | 76.0 ± 0.2 | 76.9 ± 0.2 | 0.80 ± 0.01 | 2.1 ± 0.1 | 4.5 ± 0.6 | 18.7 ± 4.1 | 4.2 ± 1.1 |
| **THF** | 76.5 ± 0.2 | 77.2 ± 0.2 | 0.75 ± 0.02 | 1.9 ± 0.1 | 1.9 ± 0.6 | 27.5 ± 2.0 (L)<br>1.39 ± 0.2 (H) | 14 ± 4.7<br>0.73 ± 0.3 |
| **NADPH** | 76.9 ± 0.4 | 75.3 ± 0.2 | 1.35 ± 0.04 | 3.6 ± 0.1 | 0.23 ± 0.01 | 8.91 ± 0.8 | 39 ± 3.9 |
| **NADPD** | 76.9 ± 0.1 | 75.9 ± 0.1 | 1.38 ± 0.03 | 3.6 ± 0.2 | 0.20 ± 0.06 | 7.48 ± 0.1 | 37 ± 11 |
| **NADP<sup>+</sup> + Folate** | 77.5 ± 0.1 | 75.8 ± 0.1 | 1.76 ± 0.04 | 4.7 ± 0.3 | / | 34.1 ± 1.5 | / |
| **NADPH + Folate** | 77.6 ± 0.3 | 75.7 ± 0.3 | 1.89 ± 0.05 | 5.0 ± 0.1 | / | 136 ± 18 | / |
| **NADPD + Folate** | 77.9 ± 0.1 | 76.0 ± 0.1 | 1.81 ± 0.06 | 4.7 ± 0.1 | / | 30.7 ± 4.1 | / |
| **NADP<sup>+</sup> + DHF** | 77.2 ± 0.5 | 75.6 ± 0.4 | 1.60 ± 0.08 | 4.5 ± 0.3 | / | 129 ± 25 | / |
| **NADPH + DHF** | 78.2 ± 0.5 | 76.4 ± 0.6 | 1.79 ± 0.07 | 4.9 ± 0.1 | / | 56.1 ± 12 | / |
| **NADPD + DHF** | 77.4 ± 0.2 | 75.5 ± 0.2 | 1.74 ± 0.15 | 4.8 ± 0.2 | / | 54.1 ± 25 | / |
| **NADP<sup>+</sup> + THF (closed)** | 76.8 ± 0.2 | 75.6 ± 0.2 | 1.62 ± 0.02 | 4.4 ± 0.1 | / | 105 ± 20 | / |
| **NADP<sup>+</sup> + THF (occluded)** | 77.2 ± 0.2 | 75.6 ± 0.2 | 1.25 ± 0.01 | 3.4 ± 0.1 | / | 175 ± 30 | / |
| **NADPH + THF (closed)** | 78.0 ± 0.1 | 76.2 ± 0.1 | 1.80 ± 0.03 | 4.6 ± 0.1 | / | 120 ± 8.7 | / |
| **NADPH + THF (occluded)** | 77.4 ± 0.1 | 76.2 ± 0.1 | 1.28 ± 0.09 | 3.0 ± 0.1 | / | 158 ± 17 | / |
| **NADPD + THF (closed)** | 77.5 ± 0.2 | 75.8 ± 0.2 | 1.80 ± 0.02 | 4.8 ± 0.1 | / | 103 ± 5.0 | / |
| **NADPD + THF (occluded)** | 76.9 ± 0.2 | 75.8 ± 0.2 | 1.23 ± 0.05 | 3.3 ± 0.1 | / | 177 ± 19 | / |

### Ternary complex formation from the closed conformation

The transition-state closed configuration was probed using folate (33.6 μM, *trans)* in combination with NADP^+^ (1.5 μM, *cis),* NADPH (26.6 μM, *cis)* or NADPD (24.2 μM, *cis).* The formation of the ternary complexes was observed as transient current enhancements from the enzyme:FOL or enzyme:NADPH level (**Fig. 3A,C**). The single-molecule nature of the nanopore experiment allowed measuring the sequential order of binding and releasing of the ligands to and from the enzyme (**Fig. 3B**). When NADPH and folate were used, the majority (86.2 ± 4.2%) of the collected events leading to the ternary complex appeared from the closed (NADPH-bound or E:NADPH) configuration. Since with the concentrations tested here both binding sites have the same probability of being occupied, these results indicate that the ternary complex is formed hierarchically from the closed to the occluded conformation. Once the Michaelis complex was formed, in 86.0 ± 4.1% of measured events folate was released before NADPH (*k_off_* = 135.8 ± 18.3 s^-1^), indicating that folate has a lower affinity for the transition-state configuration than NADPH. By contrast, when NADP^+^ and folate were sampled, in 90.3 ± 1.7% of the observed events NADP^+^ was released before folate (*k_off_* = 34.1 ± 1.5 s^-1^, **Fig. 3A,B**), indicating that the transition-state configuration of DHFR has a higher affinity for folate than for NADP^+^. Hence, moving from E:NADPH:FOL to E:NADP^+^:FOL, which mimics the progression of the chemical step, the affinity for the cofactor decreases, while the affinity of DHFR for folate increases. When NADPD was sampled, in 99.2 ± 0.1% of the events the ternary complex was formed from the NADPD level, and in 99.6 ± 0.1% of the events the dissociation produced a NADPD-bound configuration (*k_off_* = 30.7 ± 4.1 s^-1^) (**Fig. 3A,B**), indicating that substitution with deuterium in the cofactor promotes a stronger hydrogen-bond-mediated interaction of the cofactor to the closed transition-state configuration.

**Figure 3.**
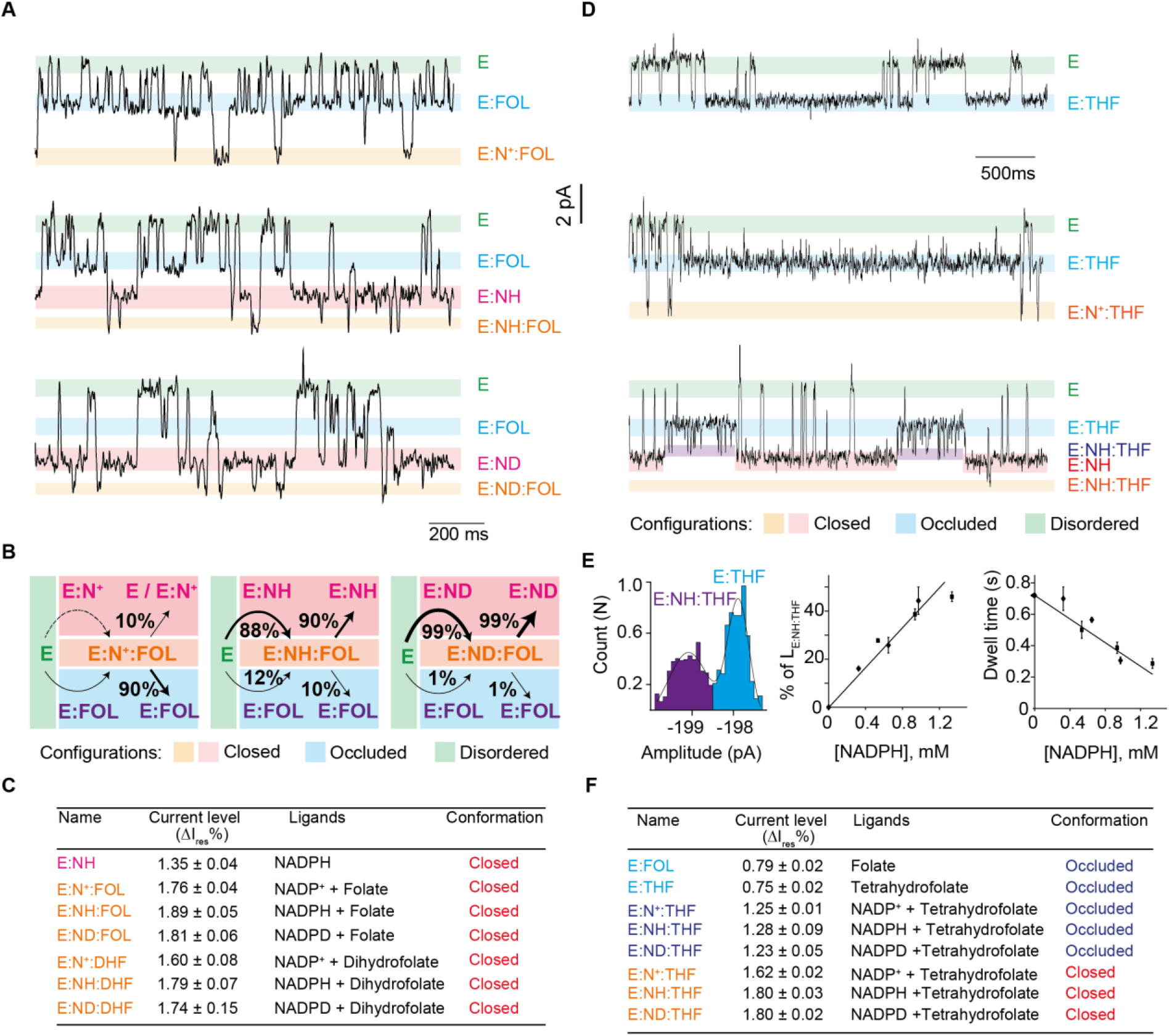
Formation of the ternary complex in the closed and occluded configuration. **(A)** Typical current blockades induced by DHFR_tag_ (50 nM, *cis*) in the presence of folate (33.6 μM, *trans)* and NADP^+^ (1.5 μM, *cis*, top), folate (33.6 μM, *trans)* and NADPH (26.6 μM, *cis,* middle) or folate (30.9 μM, *trans)* and NADPD (24.2 μM, *cis,* bottom). **(B)** Scheme indicating the hierarchy of ligand binding to DHFR_tag_ under the different conditions. **(C)** Table showing the connection between the observed current levels and the different structure of DHFR as obtained by X-ray crystallography^23^. (D) Typical tetrahydrofolate induced blockades (12.6 μM, *trans,* top), and in the presence of NADP^+^ (8.8 μM, *cis,* middle) and in the presence of NADPH (973 μM, *cis,* bottom). **(E)** Left, full point histogram of a E:NADPH:THF complex. Middle, relative occupancy of NADPH in the ternary complex (E:NADPH:THF) at different NADPH concentrations. Right, dependency of the dwell time of tetrahydrofolate *(trans)* versus the concentration of the NADPH *(cis).* The black squares are obtained when tetrahydrofolate and NADPH were used in the experiment; the black diamonds when the reaction was followed (see later). **(F)** Table showing the connection between the observed current levels and the different structure of DHFR as obtained by X-ray crystallography^23^. All current traces were collected in 250 mM KCl, 15 mM Tris-HCl pH 7.8 at room temperature (25 °C), by applying a Bessel-low pass filter with a 2 kHz cut-off and sampled at 10 kHz. The traces in were filtered digitally with a Gaussian low-pass filter with a 100 Hz cut-off.

### Ternary complex formation from the occluded conformation: product release

In the DHFR reaction, the release of the product THF is facilitated by the binding of NADPH.^39,40^ We showed earlier that THF can bind with high affinity or low affinity to the enzyme (**Fig. 2H, 3D**). In the presence of tetrahydrofolate (10 μM, *trans)* and NADPH (0.3 mM, *cis),* we observed two classes of blockades. When THF was bound to the high-affinity conformation, additional reversible current events were observed (Δ*I*_res%_ = 0.47 ± 0.01%, **Fig. 3D, Supplementary Fig. 1, 2**). Increasing the concentration of NADPH (to 0.6 or 0.9 mM) increased the frequency of the transient events but not their duration (**Fig. 3E**), indicating that the additional current enhancements reflected the transient formation of the occluded ternary product E:NADPH:THF. As observed in ensemble experiments^39^, the dwell time of the E:THF complex decreased with increasing NADPH concentration (**Fig. 3E**). Interestingly, the amplitude of the ionic current level of the occluded E:NADPH:THF ternary complex (Δ*I*_res%_ = 1.28 ± 0.09%), was similar to that of the closed E:NADPH binary complex (Δ*I*_res%_ = 1.35 ± 0.04%)(**Fig. 3C,F**). This observation is consistent with previously reported relaxation dispersion experiments^39,40^, which showed that the E:NADPH:THF complex transiently samples a closed excited state whereby the backbone conformation closely resembles the ground state of the closed E:NADPH complex.

When THF was bound to the low-affinity conformation, additional reversible NADP^+^ or NADPH current events were observed with Δ*I*_res%_ = 1.62 ± 0.02% and 1.80 ± 0.03%, respectively (**Fig. 3D,F, Supplementary Fig. 2**). These values were similar to the blockades recorded for the DHF and FOL ternary complexes, which is in the closed conformation (**Fig. 3**). It follows that the low-affinity THF events are likely to correspond to THF binding to the closed conformation of DHFR_tag_, while the high-affinity THF blockades correspond to the binding to the occluded conformation of DHFR_tag_. Interestingly, the comparison between the binding of NADP^+^ to the low- and high-affinity E:THF levels, indicated that NADP^+^ bound more often (**Fig. 3D**) and stronger (*k_off_* = 105.1 ± 20.4 s^-1^, *k_off_* = 175.0 ± 30.2 s^-1^, respectively, **Table 1, Fig. Supplementary Fig. 2**) to the low-affinity level, indicating that switching from the closed to the occluded conformation reduced the affinity of the enzyme for NADP^+^.

### Catalyzed reaction

The catalytic cycle was first sampled under physiological conditions of NADPH (0.7 mM, *cis),* DHF (4.3 μM, *trans)* and pH (7.15) (**Fig. 4A, Supplementary Fig. 3**). Multiple reactions (1.08 ± 0.04 per second) from individual enzymes as shown by the appearance of the typical THF blockades (pink asterisks, **Fig. 4A**) were observed. The reaction rate (1.08 ± 0.04 s^-1^, *N* = 3 individual nanopore experiments, *n* = 400 events) was measured by counting the number of products formed over the time of the protein blockade. The single-molecule observation revealed that not all the ternary complexes lead to a reaction. The percentage of non-reactive configurations increased with the pH (from 62.6 ± 1.9% at pH 7.15 to 89.0 ± 2.8% at pH 9.1, **Table S1, Supplementary Fig. 3-5**), suggesting that the reaction occurred if the substrate is protonated within the lifetime of the transition-state configuration (~18 ms)(**Fig. 4B**). We also found that in about 10% of the recorded events, long ternary complexes were observed (1.64 ± 1.47 sec, Blue underlining, **Fig. 4C, Table S1, Supplementary Fig. 3-5**). In the presence of NADPD the fraction of these long events was reduced by about threefold (**Table S1, Supplementary Fig. 6-8**). Since the rate-limiting hydride transfer step^22^ also shows a kinetic isotope effect of three, this suggests that the long blockades are related to the chemical step. When the reaction was sampled using a 10-fold excess of DHF compared to NADPH, the majority (58.2 ± 7.9%) of reactions still occurred from the NADPH-bound level, but only after large current rearrangements occurred (**Fig. 4D, Supplementary Fig. 9**). This confirms that the reaction is hierarchical and preferentially occurs from the closed state despite the order of binding of the substrates.

**Figure 4:**
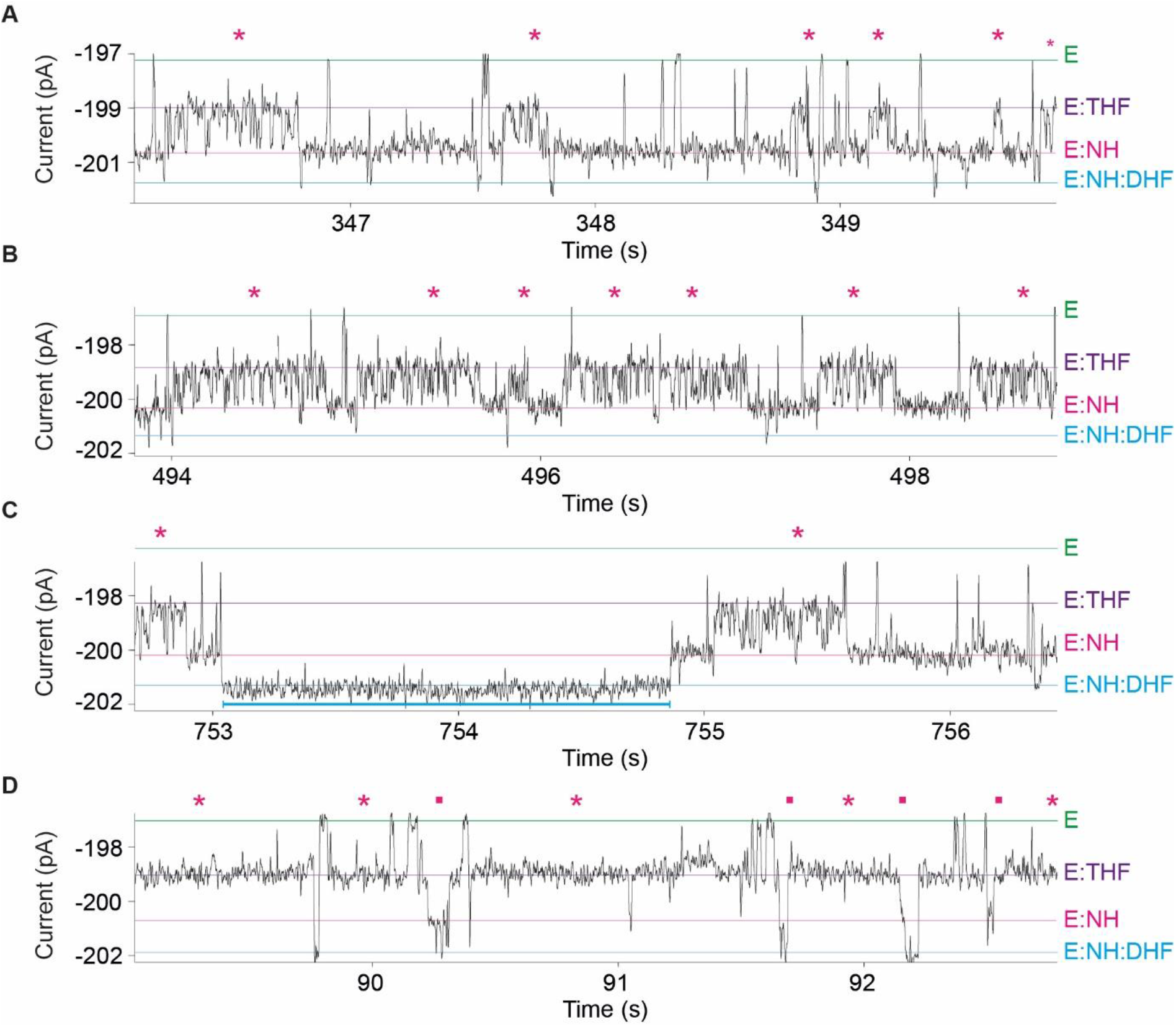
Continuous recording of an individual DHFR_tag_ enzyme catalyzed reaction. **(A-C)** The three panels are representative parts of one enzyme its catalytic trace with NADPH (0.7 mM) and DHF (4.3 μM) added to the *cis* and *trans* compartments, respectively. The pink asterisks are indicating the occurrence of the product tetrahydrofolate. **(D)** Catalytic reaction measured with an excess of DHF (50.5 μM, *trans)* compared to NADPH (5.5 μM, *cis).* The pink asterisks are indicating the occurrence of the product, tetrahydrofolate. The pink squares indicate the rearrangement in the transition-state conformation leading to the reaction. The current levels of the apo-enzyme, tetrahydrofolate-bound, NADPH-bound, and ternary complex conformations are highlighted with a green, purple, pink and blue line, respectively. The blue underlining **(C)** shows a representative long ternary complex. Current traces were collected in 250 mM KCl, 15 mM Tris-HCl at pH 7.15 **(A-C)** or 7.8 **(D)** and 25 °C, by applying a Bessel-low pass filter with a 2 kHz cut-off and sampled at 10 kHz. The trace was filtered digitally with a Gaussian low-pass filter with a 100 Hz cut-off.

## Discussion and Conclusion

It is generally assumed that enzymes fold in one conformation that stabilizes the transition state of the catalyzed reaction.^41,42^ More recently, it has emerged that dynamics are strongly entrenched in enzyme catalysis.^6,43,44^ It is now becoming evident that enzymes exist as complex ensembles of conformations that are generated by the continuous unfolding and refolding in localised regions of the protein.^45,46^ This large body of conformations differ only slightly and embody the native-state ensemble. Interconversion is fast and often on the same time-scale as the catalytic step.^47^ Although the relation between dynamics and the chemical step has been largely established^43,47,48^, what has often been overlooked is that in many enzymatic reactions, the release of product is the rate-limiting step.^22,49,50^ Hence, in such reactions an enhancement of the catalytic rate must include strategies to accelerate product release.

Notably, we previously found that DHFR exists in at least four slow-converting (hence ground-state) conformations, which were called conformers. Exchange between conformers is more frequently at the transition state^31^ or during the rate-limiting product release (**Fig. 2H**). This is in accordance with NMR relaxation experiments, which revealed that motions throughout the enzyme become activated at the transition-state configuration (E:NADP^+^:DHF)^51^. Interestingly, we found that occasionally the chemical step induces switching to a long-lasting and inactive conformer of DHFR that is not observed under non-reactive conditions *(e.g.* at high pH or with the slow-reacting folate), suggesting that conformer switching is induced by the excess enthalpy generated during the reaction. This observation suggests therefore that the soft interactions in folded enzymes are used not only to provide the ideal chemical environment for the reaction to occur, but also to dissipate the excessive energy generated during the catalytic step.

The single-molecule experiments described here allowed mapping the hierarchy of the catalytic cycle and revealed the affinity of the enzyme for the different reaction intermediates; E:NADPH, E:NADPH:DHF, E:NADP^+^:DHF, E:NADP^+^:THF, E:THF, and E:NADPH:THF. We found that the transition state is formed from the E:NADPH closed configuration rather than from the E:DHF occluded configuration (**Fig. 4D**), indicating that the binding of NADPH to DHFR induces an allosteric effect that increases the affinity of the enzyme for the substrate, positioning it in the reactive configuration. Once the transition-state conformation is obtained DHFR acquires affinity for NADP^+^, as observed by the faster release of NADP^+^ from the apo-enzyme (300 s^-1^)^22^ compared to release of NADP^+^ from the closed (E:NADP^+^:DHF) or occluded (E:NADP^+^:THF) ternary complexes of DHFR_tag_ (105 ± 20 s^-1^ and 175 ± 30 s^-1^, respectively, **Table 1**). By contrast, the other reaction product, THF, binds with low affinity to the closed configuration (off rate = 27.5 ± 2 s^-1^) and with high affinity for the occluded (off rate = 1.39 ± 0. 2 s^-1^) configuration of DHFR_tag_. Hence, the transition state has affinity for the reactants and products alike, the closed conformation has high affinity for the cofactor NADPH, the occluded conformation has high affinity for the product THF, and both ground-state conformations have low affinity for NADP^+^. Together, these evidences point towards a catalytic mechanism in which DHFR cycles between the closed and occluded conformations like a two-stroke engine. The free energy liberated in the chemical step induces a power stroke that switches the enzyme from the closed to the occluded conformation. In this step, the affinity of DHFR for the cofactor is reduced, the affinity for the product THF is increased and NADP^+^ is expelled. The free energy of the binding of NADPH to the vacated site produces a backstroke, in which DHFR switches back to the initial configuration and THF is released. Interestingly, the cycling is not always perfect. Rarely, the energy released from the chemical step produces an off-path DHFR conformations in which NADP^+^ is not efficiently expelled (**Fig. 4C**). This latter observation suggests that the network of stabilizing interactions in folded enzymes might pay a role in dissipating the energy excess arising from the chemical step.

This hierarchical conformational exchange is likely to have evolved to tightly control the affinity of the enzyme for NADP^+^. Notably, during the whole catalytic cycle, the ground-state conformations of the enzyme maintain low affinity for NADP^+^. In *E. coli* cells the concentrations of NADP^+^ and NADPH are equal. This is in contrast with human cells, where the concentration of NADPH is about 100-fold higher than NADP^+^. Although the structure of human DHFR is almost identical to that of *E. coli* DHFR, their sequence is highly divergent. Both DHFR enzymes have similar kinetics involving the same catalytic cycle with its intermediates. However, 35% of the human enzyme also populates a second catalytic cycle, with E:NADP^+^:THF being the branch point, where NADP^+^ remains bound to the enzyme and THF is first released.^52,53^ Interestingly, the Met20 loop of human DHFR remains in the closed configuration. Hence, the occluded configuration of *E. coli* DHFR, in which the formation of THF act as a thermodynamic sink, most likely evolved to promote the release of the NADP^+^, allowing efficient catalysis at high intracellular concentration of NADP^+^.

## Author information

### Author contributions

N.S.G. performed the experiments and analysed the data presented in the manuscript. N.S.G., and G.M. wrote the manuscript. G.M. supervised the project.

## Acknowledgments

The plasmid containing the alcohol dehydrogenase from *Thermoanaerobacter brockii* was a kind gift from the Allemann Group (Cardiff University).

