## Supplementary information for "Substrate binding and turnover modulate the affinity landscape of dihydrofolate reductase to increase its catalytic efficiency"

**Material and methods**

Unless otherwise specified all chemicals were bought from Sigma-Aldrich. DNA was purchased from Integrated DNA Technologies (IDT), enzymes from Fermentas and lipids from Avanti Polar Lipids. High purity (6*S*)-5,6,7,8-Tetrahydrofolic acid was purchased from Schircks Laboratories. All errors in this work are given as standard deviations.

**Sequences**

>ClyA (DNA sequence)

ATGACGGGTATCTTTGCGGAACAGACGGTGGAAGTTGTGAAAAGTGCGATTGAAACGG
CTGACGGTGCGCTGGACCTGTATAATAAATATCTGGATCAGGTCATCCCGTGGAAAACC
TTTGACGAAACGATTAAAGAACTGAGCCGTTTCAAACAGGAATACAGTCAAGAAGCGTC
CGTCCTAGTGGGCGATATCAAAGTGCTGCTGATGGATTCTCAGGACAAATATTTTGAAG
CTACCCAAACGGTTTACGAATGGGCGGGTGTGGTTACCCAGCTGCTGTCCGCATATATT
CAGCTGTTCGATGGATACAATGAGAAAAAAGCGAGCGCGCAGAAAGACATTCTGATCCG
CATTCTGGATGACGGCGTGAAAAAACTGAATGAAGCCCAGAAATCGCTGCTGACCAGCT
CTCAATCATTTAACAATGCCTCGGGTAAACTGCTGGCACTGGATAGCCAGCTGACGAAC
GACTTTTCTGAAAAAAGTTCCTATTACCAGAGCCAAGTCGATCGTATTCGTAAAGAAGCC
TACGCAGGTGCCGCAGCAGGTATTGTGGCCGGTCCGTTCGGTCTGATTATCTCATATTC
AATTGCTGCGGGCGTTGTCGAAGGTAAACTGATTCCGGAACTGAACAATCGTCTGAAAA
CCGTTCAGAACTTTTTCACCAGTCTGTCTGCTACGGTCAAACAAGCGAATAAAGATATCG
ACGCCGCAAAACTGAAACTGGCCACGGAAATCGCTGCGATTGGCGAAATCAAAACCGA
AACGGAAACCACGCGCTTTTATGTTGATTACGATGACCTGATGCTGAGCCTGCTGAAAG
GTGCCGCGAAGAAAATGATTAATACCTCTAATGAATATCAGCAGCGTCACGGTAGAAAA
ACCCTGTTTGAAGTCCCGGATGTGGGCAGCAGCTACCACCATCATCACCACTAAAAGCT
T

>ClyA-AS (Protein sequence)

MTGIFAEQTVEVVKSAIETADGALDLYNKYLDQVIPWKTFDETIKELSRFKQEYSQEASVLVG
DIKVLLMDSQDKYFEATQTVYEWAGVVTQLLSAYIQLFDGYNEKKASAQKDILIRILDDGVKK
LNEAQKSLLTSSQSFNNASGKLLALDSQLTNDFSEKSSYYQSQVDRIRKEAYAGAAAGIVAG
PFGLIISYSIAAGVVEGKLIPELNNRLKTVQNFFTSLSATVKQANKDIDAAKLKLATEIAAIGEIK
TETETTRFYVDYDDLMLSLLKGAAKKMINTSNEYQQRHGRKTLFEVPDVGSSYHHHHH

> $DHFR_{tag}$ (DNA sequence)

ATGGCTTCGGCTATGATTTCACTGATTGCGGCACTGGCTGTCGATCGTGTTATTGGTAT
GGAAAACGCTATGCCGTGGAATCTGCCGGCTGATCTGGCGTGGTTTAAACGTAACACTC

TGGACAAGCCGGTCATTATGGGCCGCCATACGTGGGAAAGCATCGGTCGTCCGCTGCC
GGGTCGCAAAAATATTATCCTGAGCAGCCAGCCGGGCGAAGATGACGAAGTGACGTGG
GTTAAGAGCGTCGATGAAGCAATTGCGGCGGCAGGCGACGTGCCGGAAATTATGGTTA
TCGGCGGTGGCCGCGTTTATGAACAGTTCCTGCCGAAAGCCCAAAAGCTGTACCTGAC
CCATATCGATGCAGAAGTCGAAGGTGATACGCACTTTCCGGACTATGAACCGGATGACT
GGGAAAGTGTGTTCTCCGAATTTCACGACGCCGACGCTCAGAACAGCCACTCATACTCA
TTCGAAATCCTGGAACGCCGTGGCAGCAGTACTCGAGCGAAAAAGAAGATTAAGAAAAA
GCTAAAACAGGGCAGCGCGTGGAGCCATCCGCAGTTTGAAAAATGATAAGCTT

> $DHFR_{tag}$ (Protein sequence)
MASAMISLIAALAVDRVIGMENAMPWNLPADLAWFKRNTLDKPVIMGRHTWESIGRPLPGR
KNIILSSQPGEDDEVTWVKSVDEAIAAAGDVPEIMVIGGGRVYEQFLPKAQKLYLTHIDAEVE
GDTHFPDYEPDDWESVFSEFHDADAQNSHSYSFEILERRGSSTRAKKKIKKKLKQGSAWSH
PQFEK

> Alcohol dehydrogenase from *Thermoanaerobacter brockii* (DNA sequence)
ATGAAAGGCTTTGCCATGCTGTCAATTGGCAAAGTTGGTTGGATTGAAAAGGAAAAGCC
GGCCCCTGGCCCGTTTGATGCAATTGTGCGCCCGCTCGCTGTGGCACCATGTACGTCG
GATATTCATACCGTGTTCGAAGGCGCCATTGGGGAACGCCACAACATGATTCTGGGACA
TGAAGCCGTAGGCGAAGTGGTAGAAGTGGGTAGCGAAGTCAAAGACTTCAAGCCCGGA
GACCGCGTCGTTGTACCGGCAATTACCCCGGATTGGCGTACCAGCGAAGTCCAGCGTG
GATACCATCAGCACAGCGGCGGCATGCTGGCGGGTTGGAAATTTAGCAATGTGAAAGA
CGGCGTTTTCGGTGAGTTTTTTCATGTAAACGATGCAGATATGAACCTGGCACATCTGC
CGAAAGAAATTCCGTTGGAAGCTGCCGTGATGATTCCAGATATGATGACAACAGGTTTC
CACGGGGCGGAACTGGCGGATATCGAATTAGGGGCGACGGTAGCCGTGCTGGGCATT
GGCCCTGTTGGGCTGATGGCTGTCGCCGGTGCAAAGCTGCGCGGTGCGGGACGTATC
ATTGCGGTGGGTAGTCGTCCGGTCTGCGTAGATGCGGCCAAGTATTATGGCGCGACCG
ATATTGTTAATTACAAGGACGGCCCGATCGAGAGCCAGATCATGAATCTGACCGAAGGG
AAAGGTGTGGATGCCGCGATTATCGCCGGAGGGAACGCGGATATTATGGCGACCGCG
GTCAAGATCGTCAAGCCCGGGGGCACGATCGCCAACGTTAATTATTTCGGTGAAGGCG
AGGTTCTGCCTGTACCGCGTCTTGAGTGGGGTTGCGGCATGGCACATAAAACCATTAAA
GGTGGCCTGTGTCCTGGAGGGCGCTTACGCATGGAACGCCTTATCGATCTGGTGTTTT
ATAAACGTGTGGACCCGTCCAAACTGGTTACCCATGTATTTCGCGGCTTTGATAACATTG
AGAAAGCGTTTATGCTGATGAAAGATAAGCCAAAAGATCTGATCAAACCAGTCGTGATCT
TGGCG

> Alcohol dehydrogenase from *Thermoanaerobacter brockii* (protein sequence)
MKGFAMLSIGKVGWIEKEKPAPGPFDAIVRPLAVAPCTSDIHTVFEGAIGERHNMILGHEAVG

EVVEVGSEVKDFKPGDRVVVPAITPDWRTSEVQRGYHQHSGGMLAGWKFSNVKDGVFGE
FFHVNDADMNLAHLPKEIPLEAAVMIPDMMTTGFHGAELADIELGATVAVLGIGPVGLMAVA
GAKLRGAGRIIAVGSRPVCVDAAKYYGATDIVNYKDGPIESQIMNLTEGKGVDAAIIAGGNAD
IMATAVKIVKPGGTIANVNYFGEGEVLPVPRLEWGCGMAHKTIKGGLCPGGRLRMERLIDLV
FYKRVDPSKLVTHVFRGFDNIEKAFMLMKDKPKDLIKPVVILA

**His-tagged Type I ClyA-AS protein overexpression and purification**

*E. cloni*® EXPRESS BL21 (DE3) cells were transformed with the pT7-SC1 plasmid containing the ClyA-AS gene. ClyA-AS contains eight mutations relative to the *S. Typhi* ClyA-WT: C87A, L99Q, E103G, F166Y, I203V, C285S, K294R and H307Y (the H307Y mutation is in the C-terminal hexahistidine-tag added for purification).[33] Transformants were selected after overnight growth at 37 ºC on LB agar plates supplemented with 100 µg/mL ampicillin. The resulting colonies were grown at 37 ˚C (200 rpm shaking) in 2xYT medium supplemented with 100 µg/mL ampicillin until the O.D. at 600 nm was ~0.8. ClyA-AS expression was then induced by addition of 0.5 mM IPTG, and the temperature was switched to 25 °C for overnight growth (200 rpm shaking). The next day the bacteria were harvested by centrifugation at 6000x*g* for 25 min at 4 °C and the pellets were stored at -80 ºC.

Pellets containing monomeric ClyA-AS arising from 50 mL culture were thawed and resuspended in 20 mL of wash buffer (10 mM imidazole, 150 mM NaCl, 15 mM Tris-HCl pH 7.5), supplemented with 1 mM $MgCl_2$ and 0.2 units/mL of DNaseI. After lysis of the bacteria by probe sonication, the crude lysates were clarified by centrifugation at 6000x*g* for 20 min at 4 °C and the supernatant was mixed with 200 µL of Ni-NTA resin (Qiagen) equilibrated in wash buffer. After 1 hr, the resin was loaded into a column (Micro Bio Spin, Bio-Rad) and washed with ~5 mL of the wash buffer. ClyA-AS was eluted with approximately ~0.5 mL of wash buffer containing 300 mM imidazole. ClyA-AS monomers were stored at 4˚C until further use.

ClyA-AS monomers were oligomerized by addition of 0.5% β-dodecylmaltoside (DDM, GLYCON Biochemicals, GmbH) and incubation at 37 ºC for 30 min. ClyA-AS oligomers were separated from monomers by blue native polyacrylamide gel electrophoresis (BN-PAGE, Bio-rad) using 4-20% polyacrylamide gels. The bands corresponding to Type I ClyA-AS were excised from the gel and placed in 150 mM NaCl, 15 mM Tris-HCl pH 7.5, supplemented with 0.2% DDM and 10 mM EDTA to allow diffusion of the proteins out of the gel.

**Overexpression of the $DHFR_{tag}$ protein**

The pT7-SC1 plasmid containing the $DHFR_{tag}$ gene[30] was transformed into *E. cloni*® EXPRESS BL21(DE3) cells (Lucigen) by electroporation, and transformants were selected on LB agar plates supplemented with 100 µg/mL ampicillin and 1% glucose after overnight growth at 37 ˚C. The resulting colonies were grown at 37 ˚C in 2xYT medium supplemented with 100 µg/mL ampicillin until the optical density at 600 nm ($OD_{600}$) reached ~0.8 (200 rpm shaking). The $DHFR_{tag}$ expression was subsequently induced by addition of 0.5 mM IPTG (isopropyl β-D-1-thiogalactopyranoside), and the temperature was switched to 25 ºC for overnight growth (200 rpm shaking). The next day the bacteria were harvested by centrifugation at 6000x*g* at 4 ºC for 25 min and the resulting pellets were frozen at -80 ºC until further use.

**Purification of the $DHFR_{tag}$ protein**

Bacterial pellets originating from 50 mL culture were resuspended in 30 mL lysis buffer (150 mM NaCl, 15 mM Tris-HCl pH 7.5, 1 mM $MgCl_2$, 0.2 units/mL DNaseI, 10 µg/mL lysozyme) and incubated at 37 ºC for 20 min. After further disruption of the bacteria by probe sonication the crude lysate was clarified by centrifugation at 6000x*g* at 4 ˚C for 30 min. The supernatant was allowed to bind to ~150 µL (bead volume) of Strep-Tactin® Sepharose® (IBA) pre-equilibrated with the wash buffer (150 mM NaCl, 15 mM Tris-HCl pH 7.5) - "end over end" mixing. The resin was then loaded onto a column (Micro Bio Spin, Bio-Rad) and washed with ~20 column volumes of the wash buffer. Elution of $DHFR_{tag}$ from the column was achieved by addition of ~100 µL of elution buffer (150 mM NaCl, 15 mM Tris-HCl pH 7.5, ~15 mM D-Desthiobiotin (IBA)). Proteins were aliquoted and frozen at -20 ºC until further use. New aliquots of $DHFR_{tag}$ were thawed prior to every experiment.

**Purification of *Thermoanaerobacter brockii* alcohol dehydrogenase (TbADH)**

A pET11C plasmid containing the TbADH gene[54] was transformed into *E. cloni*® EXPRESS BL21(DE3) cells (Lucigen) by electroporation and transformants were selected on LB agar plates supplemented with 100 µg/ml ampicillin and 1% glucose after overnight growth at 37 ˚C. The resulting colonies were grown at 37 ˚C in 2xYT medium supplemented with 100 µg/mL ampicillin until the optical density at 600 nm ($OD_{600}$) reached ~0.8 (200 rpm shaking). The expression of TbADH was induced by addition of 0.5 mM IPTG (isopropyl β-D-1-thiogalactopyranoside). The cell cultures were further grown overnight at 25 °C. The next day the bacteria were harvested by centrifugation at 6000x*g* at 4 °C for 25 minutes and the pellets were stored at -80 °C until further use. The pellets containing TbADH, originating from 50 mL culture were resuspended in 10 mL of lysis buffer (25 mM Tris-HCl, pH 7.3, 0.1 mM DDT, 0.1 EDTA, 0.2 units/ml DNaseI and 10 µg/ml lysozyme) and incubated for 30 min at 37 °C. The bacteria were then lysed by 6 cycle of probe sonication (15 sweeps at 50-70% output power followed by 30 s on ice) (Branson). The clarified lysate was incubated at 80 °C for 30 minutes to precipitate *E.coli* proteins. The clarified solution was purified by anion-exchange chromatography on diethyl-aminoethyl resin (GE Healthcare) eluting with a 0-1M NaCl gradient to yield ± 65% pure TbADH as judged by SDS-PAGE. The TbADH was freeze-dried and stored at -80 °C until further use.

**Synthesis of dihydrofolate ($FH_2$)**

Ascorbic acid (1 g) was dissolved in 10 mL $H_2O$ and the pH was slowly adjusted to 6.0 with 1M NaOH. Folic acid (38.2 mg) was dissolved in 1.6 mL of 0.1 M NaOH. Both solution where mixed while stirring and while purging with $N_2$. Sodium dithionite was then added (400 mg) until dissolved and the reaction was cooled to 4 °C on ice. Under constant stirring, the pH was slowly taken to 2.8 *via* drop-wise addition of 1 M HCl with a syringe-pushing machine at a rate of 0.1/min (± 60 minutes). The precipitate was decanted and centrifuged at 4 °C, 6500x*g* for 5 minutes. The precipitate (solid pellet) was resuspended in 10 mL ice-cold 10% Sodium ascorbate (0.55 M) pH 6.0 solution. The dissolved dihydrofolate was re-crystallised by decreasing the pH to 2.8 *via* drop-wise addition of 1M HCl with a syringe pushing machine at a rate of 0.1/min (± 60 minutes). The precipitate was decanted and centrifuged at 4 °C, 6500x*g* for 5 minutes. The pellet was washed three times with 5 mL 0.001 M HCl. The product was finally resuspended in 0.001 M HCl, aliquoted and stored at -80 °C until further use. New aliquots of dihydrofolate were thawed and dissolved in 1M NaOH prior to every experiment.[55]

**Enzymatic preparation of 4R-$^2$H-nicotinamide adenine dinucleotide phosphate reduced form (NADPD)**

NADP$^+$ (10 mg) and freeze-dried TbADH (1 mg) were dissolved in 2 mL of a 25 mM Tris-HCl solution (pH 9.0) containing 0.2 mL 2-Propanol-$d_8$. The reaction mixture was incubated at 40 °C while shaking for 30 minutes. Synthesised NADPD was then purified by using anion-exchange chromatography on a HiTrap Q HP column (GE Healthcare) eluting with a 0-1M NaCl gradient (pH 7.8) as previously shown[56]. The absorbance of the eluted fraction at 260 and 340 nm was recorded. Fraction with $A_{260}/A_{340}$ ratio of < 2.4 stored at -80 °C until further use.

**Electrical recordings in planar lipid bilayers**

By convention the applied potential refers to the potential of the *trans* electrode in the planar lipid bilayer set up. ClyA-AS nanopores were inserted into lipid bilayers from the *cis* compartment, which is connected to the ground electrode. The setup consisted of two chambers separated by a 25 µm thick polytetrafluoroethylene film (Goodfellow Cambridge Limited) containing an aperture of approximately 100 µm in diameter. The aperture was pre-treated with a solution of hexadecane [10% (v/v) in pentane] to leave a hydrophobic coating to support bilayer formation. 1,2-diphytanoyl-*sn*-glycero-3-phosphocholine (DPhPC) in pentane (10 mg/mL) was then added to the buffer (500 µL, 250 mM KCl, 15 mM Tris-HCl, pH 7.8) present in both compartments. After evaporation of the pentane, a lipid monolayer at the air–water interface can self-assemble spontaneously by lowering and raising the buffer across the aperture. 0.01–0.1 ng of purified ClyA-AS were added to the grounded *cis* compartment. The reconstitution of single nanopores was monitored electrically by applying a -35 mV bias. ClyA-AS nanopores display a higher open pore current at positive than at negative applied potentials, which provides a useful tool to determine the orientation of the pore. Electrical recordings were carried out in 250 mM KCl, 15 mM Tris-HCl pH 7.8.

**Data recordings and analysis**

Electrical signals from planar lipid bilayer recordings were amplified using an Axopatch 200B patch clamp amplifier (Axon Instruments) and digitized with a Digidata 1440 A/D converter (Axon Instruments). Data were recorded using the Clampex 10.5 software (Molecular Devices) and the subsequent analysis was carried out with the Clampfit software (Molecular Devices). Electrical recordings were carried out in 250 mM KCl, 15 mM Tris-HCl pH 7.8., by applying a 2 kHz low-pass Bessel filter and a 10 kHz sampling rate.

Residual current values ($I_{res\%}$) of the different $DHFR_{tag}$ variants were calculated by $I_{res\%} = I_B/I_o$, in which $I_B$ and $I_o$ represent the blocked an open-pore current values, respectively. $I_B$ and $I_o$ values were calculated from Gaussian fits to all point current histograms (0.1 pA bin size) from at least 3 individual single channels each displaying at least 50 current blockades. For analysis of substrate binding events to $DHFR_{tag}$, traces were filtered digitally with a Bessel (Gaussian) low-pass filter with a 100 Hz cut-off. Current transitions from *apo* to substrate:bound were analysed with the 'single channel search' option in Clampfit. Events shorter than 0.1 ms were ignored. The process of event collection was monitored manually. The resulting event dwell times ($\tau_{off}$) and the times between events ($\tau_{on}$) were binned together as cumulative distributions and fitted to a single exponential to retrieve the ligand-induced lifetimes $\tau$ ($\tau_{off}$) and the ligand-induced inter-event times ($\tau_{on}$). The average amplitude of the events was derived from Gaussian fits to the conventional distributions of the events´ amplitudes. Values for $k_{off}$ were determined as $1/\tau_{off}$, and the values for $k_{on}$ were determined by $k_{on} = 1/(\tau_{on}*[C])$ with [C] the concentration of ligand added to the *cis* or *trans* solution. Final values for $\tau_{on}$, $\tau_{off}$, $k_{on}$, $k_{off}$ and event amplitudes are the averages derived from at least three single channel experiments, each analysing at least 100 binding events on more than five different $DHFR_{tag}$ blockades. Reactant concentration were determined spectrophotometrically using extinction coefficients of 6200 $cm^{-1}$ $M^{-1}$ at 339 nm for NADPH and NADPD[22] , 22100 $cm^{-1}$ $M^{-1}$ at 302 nm for MTX[57],

28000 $cm^{-1}$ $M^{-1}$ at 297 nm for tetrahydrofolate[58], and 27000 $cm^{-1}$ $M^{-1}$ at 282 nm for folate[59] and 28000 $cm^{-1}$ $M^{-1}$ at 282 dihydrofolate[59]. Graphs were made with Origin (OriginLab Corporation). Errors are shown as standard deviations.

**Supplementary information**

**Table S1:** Table with amount of product and complexes formed per second, percentage of unreactive complexes, different dwell times for the ternary complex (τ in ms) and the distribution of occurrence of those two complexes in percentage for the different conditions. In all cases 0.6 mM cofactor was added to *cis* (NADPH or NADPD) and 2.5 μM of dihydrofolate (DHF) in *trans*. All current traces were collected in 250 mM KCl, 15 mM Tris-HCl at room temperature (25 °C) at -80 mV.

| | Reactions per second | Complex per second | Non-reactive complexes (%) | Dwell time ($\tau_1$)(ms) | Distributation ($\tau_1$)(%) | Dwell time ($\tau_2$)(s) | Distributation ($\tau_2$)(%) |
|---|---|---|---|---|---|---|---|
| NADPH pH 7.15 | 1.08 ± 0.04 | 1.88 ± 0.21 | 62.6 ± 1.9 | 22.9 ± 15.7 | 89.6 ± 9.1 | 1.2 ± 0.4 | 10.4 ± 9.1 |
| NADPH pH 7.8 | 0.68 ± 0.15 | 2.28 ± 0.44 | 76.5 ± 1.4 | 18.4 ± 4.0 | 90.3 ± 3.7 | 1.6 ± 1.5 | 9.7 ± 3.7 |
| NADPH pH 9.1 | 0.09 ± 0.02 | 2.59 ± 0.73 | 89.0 ± 2.8 | 27.7 ± 13.6 | 72.3 ± 13.6 | 0.7 ± 0.1 | 16.9 ± 13.6 |
| NADPD pH 7.15 | 0.63 ± 0.02 | 2.23 ± 0.06 | 78.1 ± 1.1 | 21.8 ± 20.8 | 98.0 ± 2.4 | 0.7 ± 0.6 | 2.0 ± 2.4 |
| NADPD pH 7.8 | 0.33 ± 0.12 | 1.72 ± 0.54 | 83.8 ± 2.8 | 27.3 ± 14.4 | 93.6 ± 3.8 | 1.1 ± 0.5 | 6.4 ± 3.8 |
| NADPD pH 9.1 | 0.04 ± 0.02 | 2.67 ± 0.61 | 94.5 ± 2.7 | 17.9 ± 6.2 | 94.0 ± 2.1 | 1.1 ± 0.8 | 6.0 ± 2.1 |

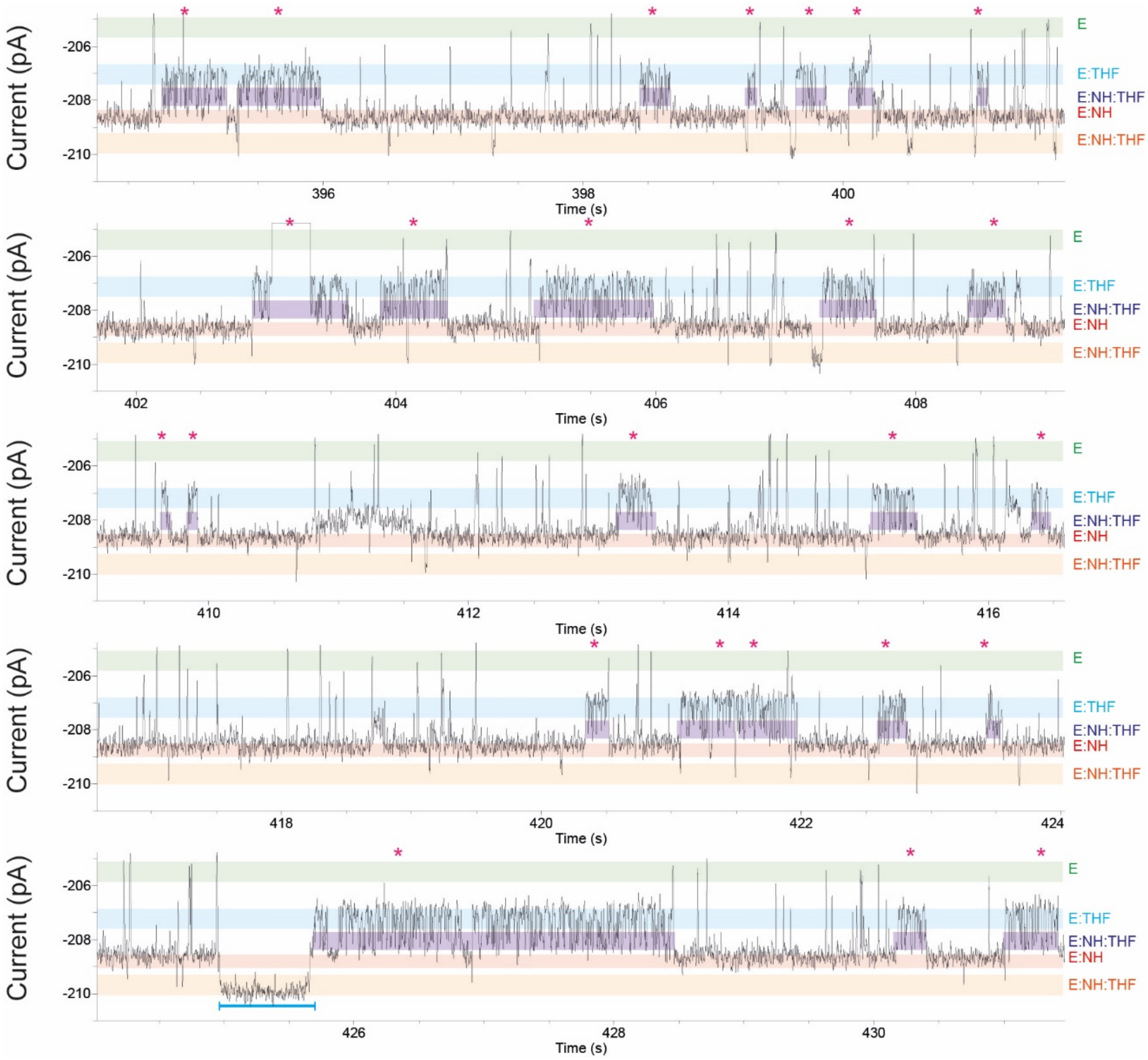


**Figure S1: Continuous recording of DHFR$_{tag}$ conformational changes during the binding of THF and NADPH.** NADPH (973 µM) and tetrahydrofolate (THF) (10 µM) were added to the *cis* and *trans* chambers, respectively. The pink asterisks are indicating the binding of tetrahydrofolate with NADPH. The current level of the *apo*-enzyme, tetrahydrofolate-bound level, NADPH-tetrahydrofolate-bound level, NADPH-bound and ternary complex level are shown as green, blue, purple, red and orange line, respectively. Current traces were collected in 250 mM KCl, 15 mM Tris-HCl pH 7.8 at room temperature (25 °C), by applying a Bessel-low pass filter with a 2 kHz cut-off and sampled at 10 kHz. The trace was filtered digitally with a Gaussian low-pass filter with a 100 Hz cut-off.

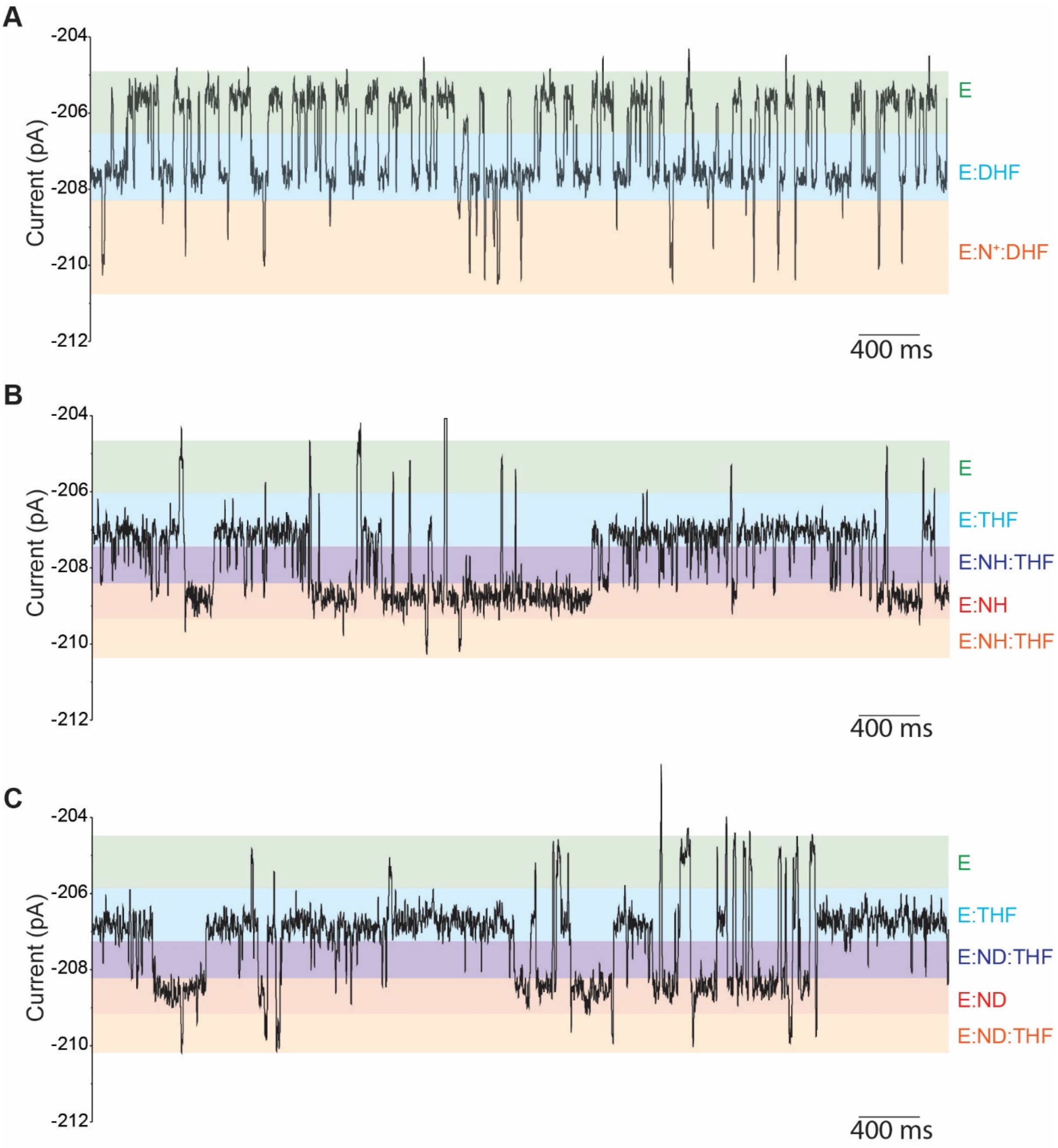


**Figure S2: Binding of the ternary complex with dihydrofolate or tetrahydrofolate. (A)** Typical current blockades induced by $DHFR_{tag}$ (~50 nM, *cis*), in the presence of $NADP^+$ (3.2 μM, *cis*) and dihydrofolate (5 μM, *trans*). **(B)** Typical current blockades induced by $DHFR_{tag}$ (~50 nM, *cis*), in the presence of NADPH (324 μM, *cis*) and tetrahydrofolate (10 μM, *trans*). **(C)** Typical current blockades induced by $DHFR_{tag}$ (~50 nM, *cis*), in the presence of NADPD (141 μM, *cis*) and tetrahydrofolate (27.3 μM, *trans*). The current level of the *apo*-enzyme, dihydrofolate or tetrahydrofolate-bound level, NADPH- or NADPD-tetrahydrofolate-bound level, NADPH or NADPD-bound and ternary complex level are shown as green, blue, purple, red and orange line, respectively. All current traces were collected in 250 mM KCl, 15 mM Tris-HCl pH 7.8 at room temperature (25 °C), by applying a Bessel-low pass filter with a 2 kHz cut-off and sampled at 10 kHz. The traces in were filtered digitally with a Gaussian low-pass filter with a 100 Hz cut-off.

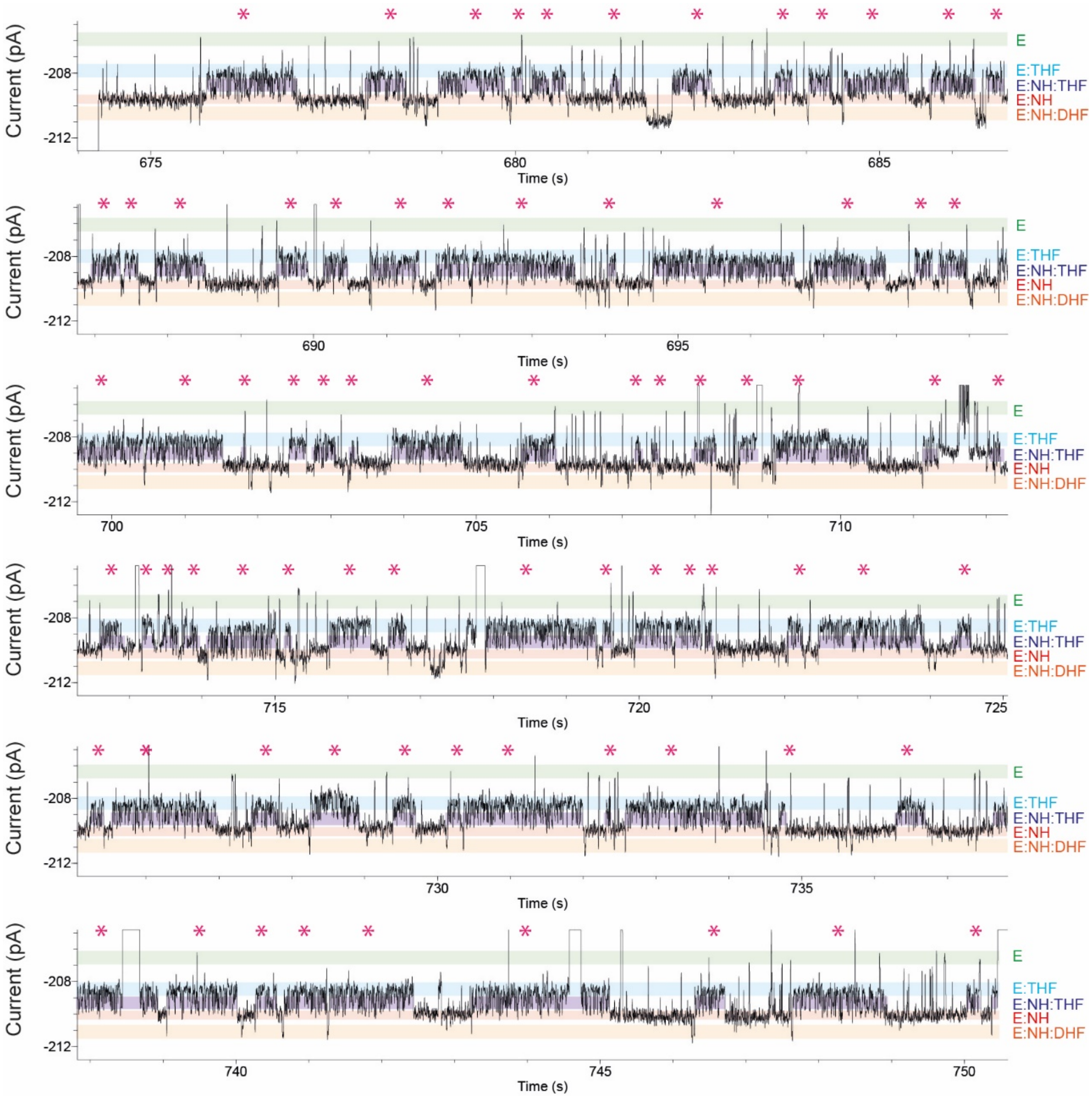


**Figure S3: Continuous recording of DHFR conformational changes during the catalysed reactions at pH 7.15.** NADPH (0.7 mM) and dihydrofolate (DHF) (4.3 μM) were added to the *cis* and *trans* chambers, respectively. The pink asterisks show the occurrence of the product, tetrahydrofolate. The current level of the *apo*-enzyme, tetrahydrofolate-bound level, NADPH-tetrahydrofolate-bound level, NADPH-bound and ternary complex level are shown as green, blue, purple, red and orange line, respectively. Current traces were collected in 250 mM KCl, 15 mM Tris-HCl pH 7.15 at room temperature (25 °C), by applying a Bessel-low pass filter with a 2 kHz cut-off and sampled at 10 kHz. The trace was filtered digitally with a Gaussian low-pass filter with a 100 Hz cut-off.

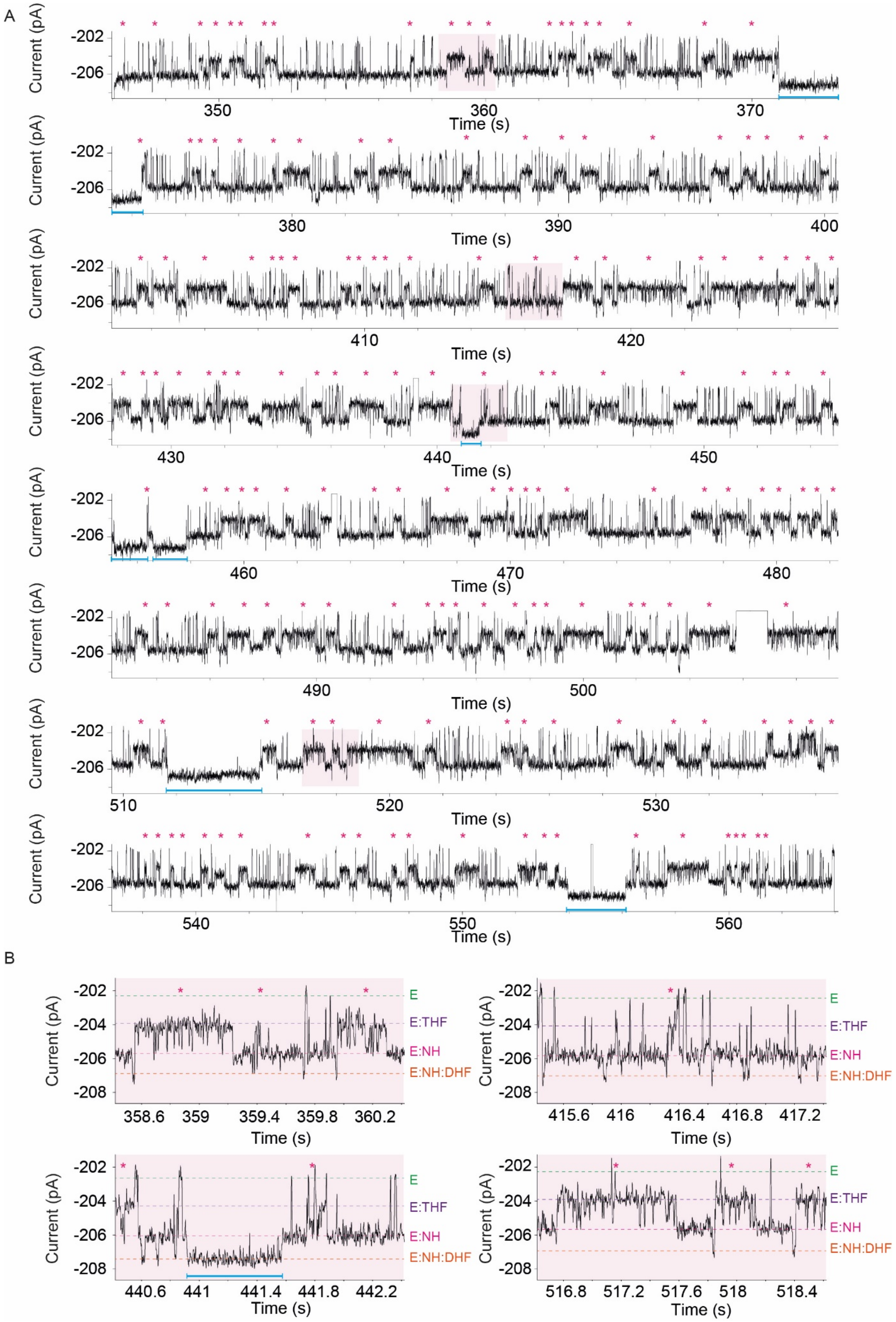


**Figure S4: Continuous recording of DHFR$_{tag}$ conformational changes during the catalysed reactions. (A)** NADPH (1.0 mM) and dihydrofolate (DHF) (1.9 µM) were added to the *cis* and *trans* chambers, respectively. The pink asterisks are indicating the occurrence of the product, tetrahydrofolate. **(B)** Selected expansions of panel A showing the details of the current levels of the catalytic reaction. The current level of the *apo*-enzyme, tetrahydrofolate-bound level, NADPH-bound level, and ternary complex level are shown as green, purple, pink and orange line, respectively. Current traces were collected in 250 mM KCl, 15 mM Tris-HCl pH 7.8 at room temperature (25 °C), by applying a Bessel-low pass filter with a 2 kHz cut-off and sampled at 10 kHz. The trace was filtered digitally with a Gaussian low-pass filter with a 100 Hz cut-off.

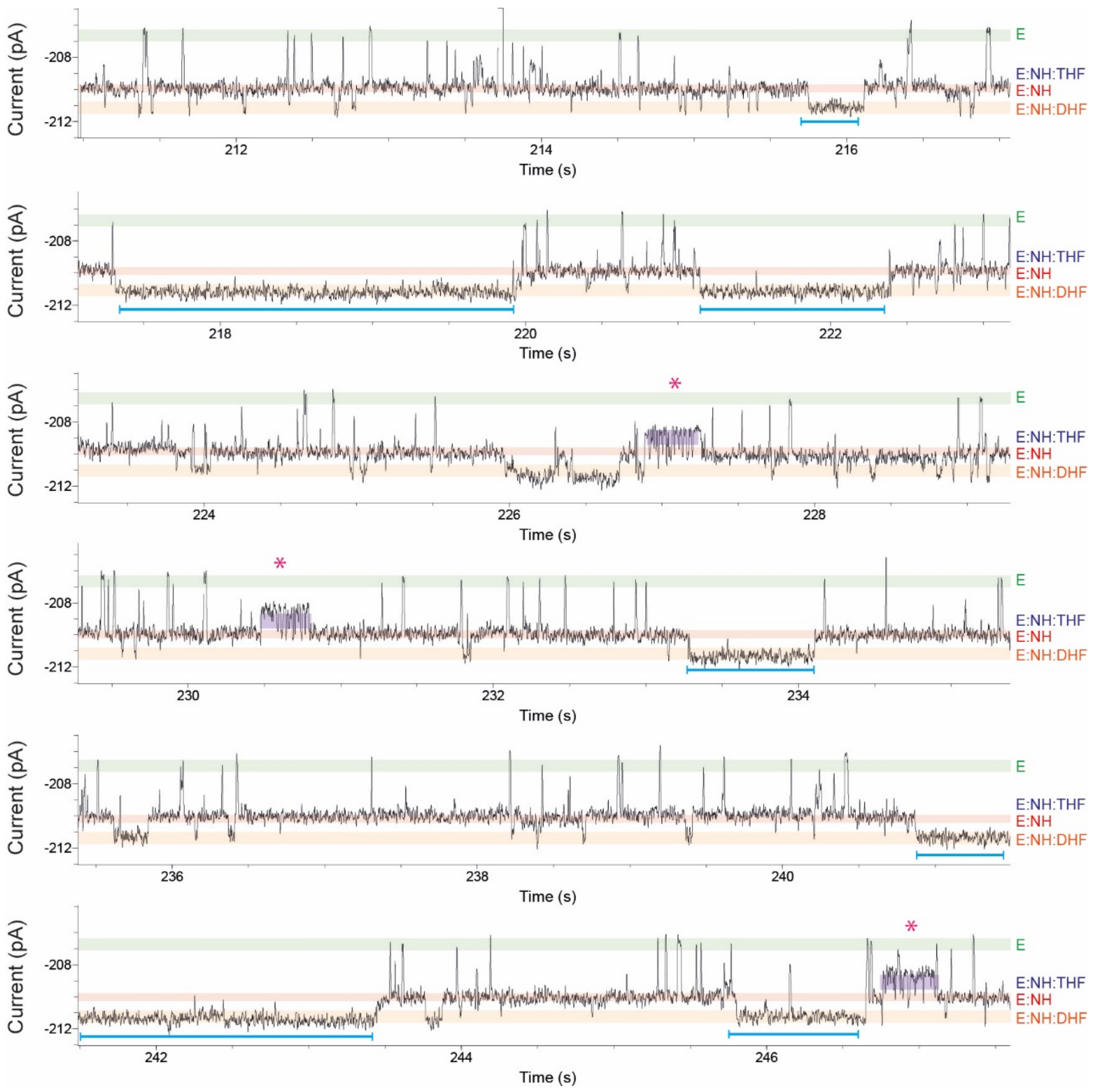


**Figure S5: Continuous recording of DHFR$_{tag}$ conformational changes during the catalysed reactions at pH 9.1.** NADPH (0.6 mM) and dihydrofolate (DHF) (1.2 μM) were added to the *cis* and *trans* chambers, respectively. The current level of the *apo*-enzyme, NADPH-tetrahydrofolate-bound level, NADPH-bound and ternary complex level are shown as green, purple, red and orange line, respectively. The pink asterisks are indicating the occurrence of the product, tetrahydrofolate. Current traces were collected in 250 mM KCl, 15 mM Tris-HCl pH 9.1 at room temperature (25 °C), by applying a Bessel-low pass filter with a 2 kHz cut-off and sampled at 10 kHz. The trace was filtered digitally with a Gaussian low-pass filter with a 100 Hz cut-off.

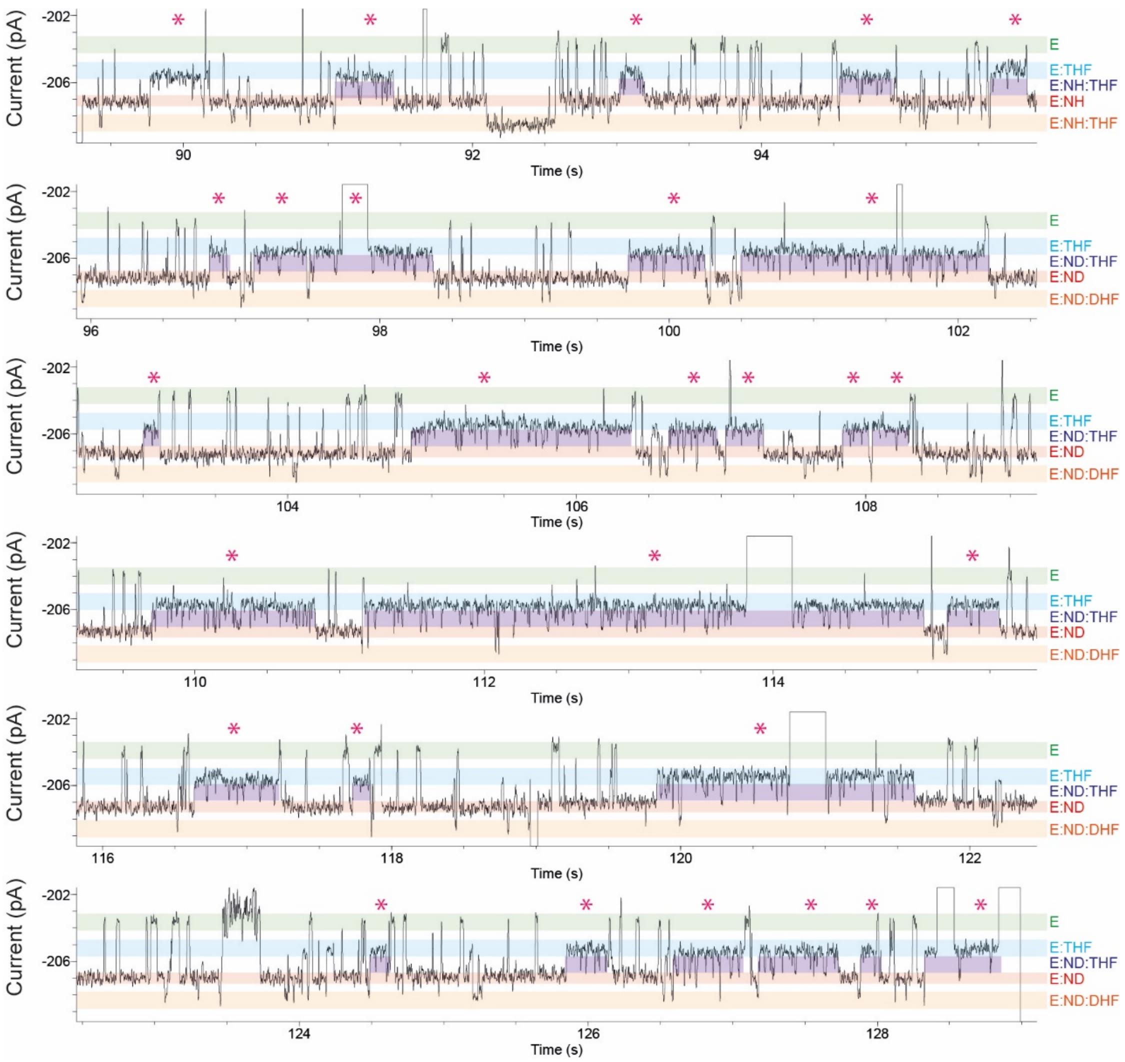


**Figure S6: Continuous recording of $DHFR_{tag}$ conformational changes during the catalysed reactions at with deuterated cofactor at low pH 7.15.** NADPD (0.7 mM) and dihydrofolate (DHF) (2.1 µM) were added to the *cis* and *trans* chambers, respectively. The pink asterisks are indicating the occurrence of the product, tetrahydrofolate. The current level of the *apo*-enzyme, tetrahydrofolate-bound level, NADPD-tetrahydrofolate-bound level, NADPD-bound and ternary complex level are shown as green, blue, purple, red and orange line, respectively. Current traces were collected in 250 mM KCl, 15 mM Tris-HCl pH 7.15 at room temperature (25 °C), by applying a Bessel-low pass filter with a 2 kHz cut-off and sampled at 10 kHz. The trace was filtered digitally with a Gaussian low-pass filter with a 100 Hz cut-off.

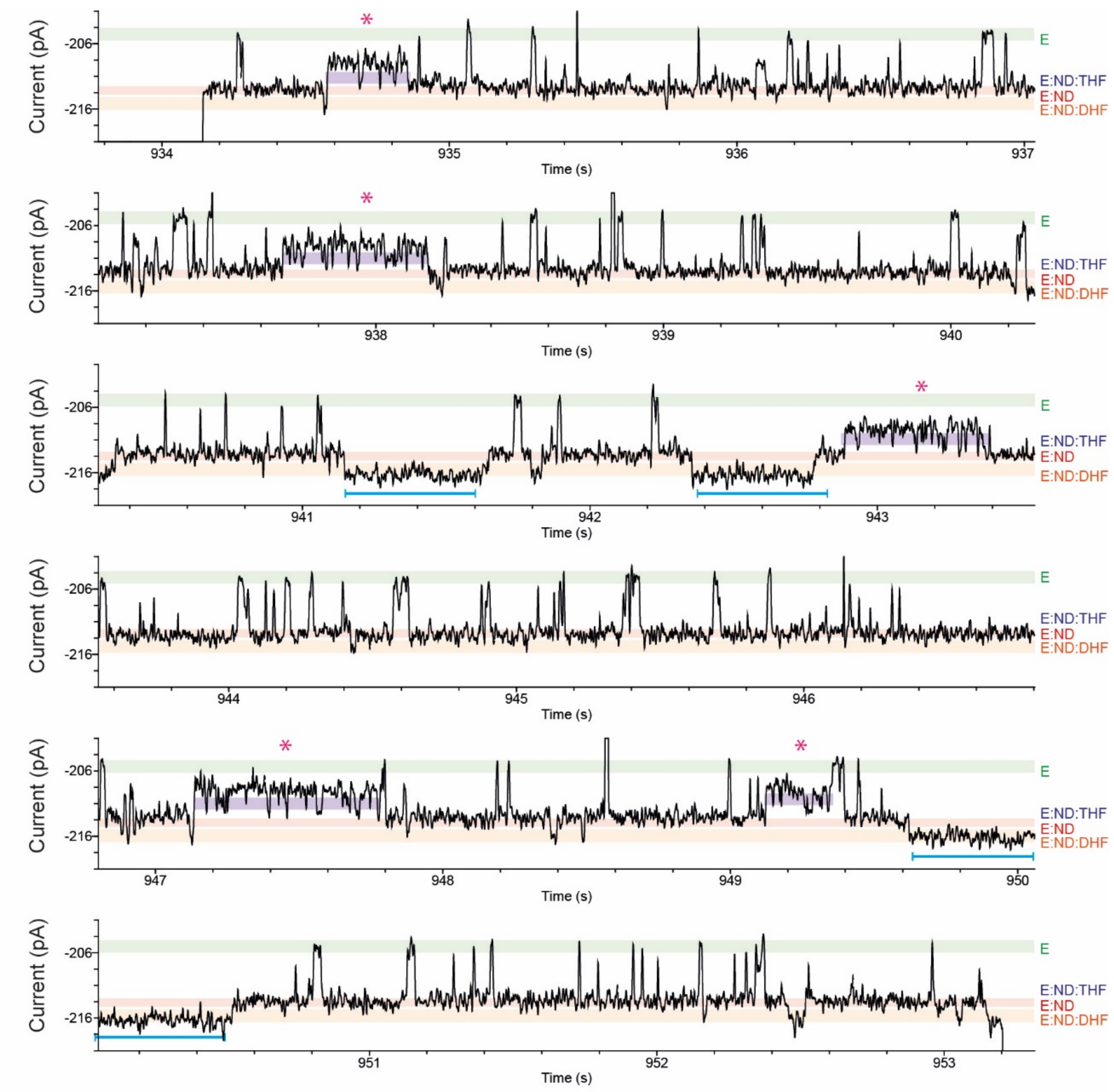


**Figure S7: Continuous recording of DHFR$_{tag}$ conformational changes during the catalysed reactions with deuterated cofactor.** NADPD (0.7 mM) and dihydrofolate (DHF) (1.3 µM) were added to the *cis* and *trans* chambers, respectively. The pink asterisks are indicating the occurrence of the product, tetrahydrofolate. The current level of the *apo*-enzyme, NADPD-tetrahydrofolate-bound level, NADPD-level and ternary complex level are shown as green, purple, red and orange line, respectively. Current traces were collected in 250 mM KCl, 15 mM Tris-HCl pH 7.8 at room temperature (25 °C), by applying a Bessel-low pass filter with a 2 kHz cut-off and sampled at 10 kHz. The trace was filtered digitally with a Gaussian low-pass filter with a 100 Hz cut-off.

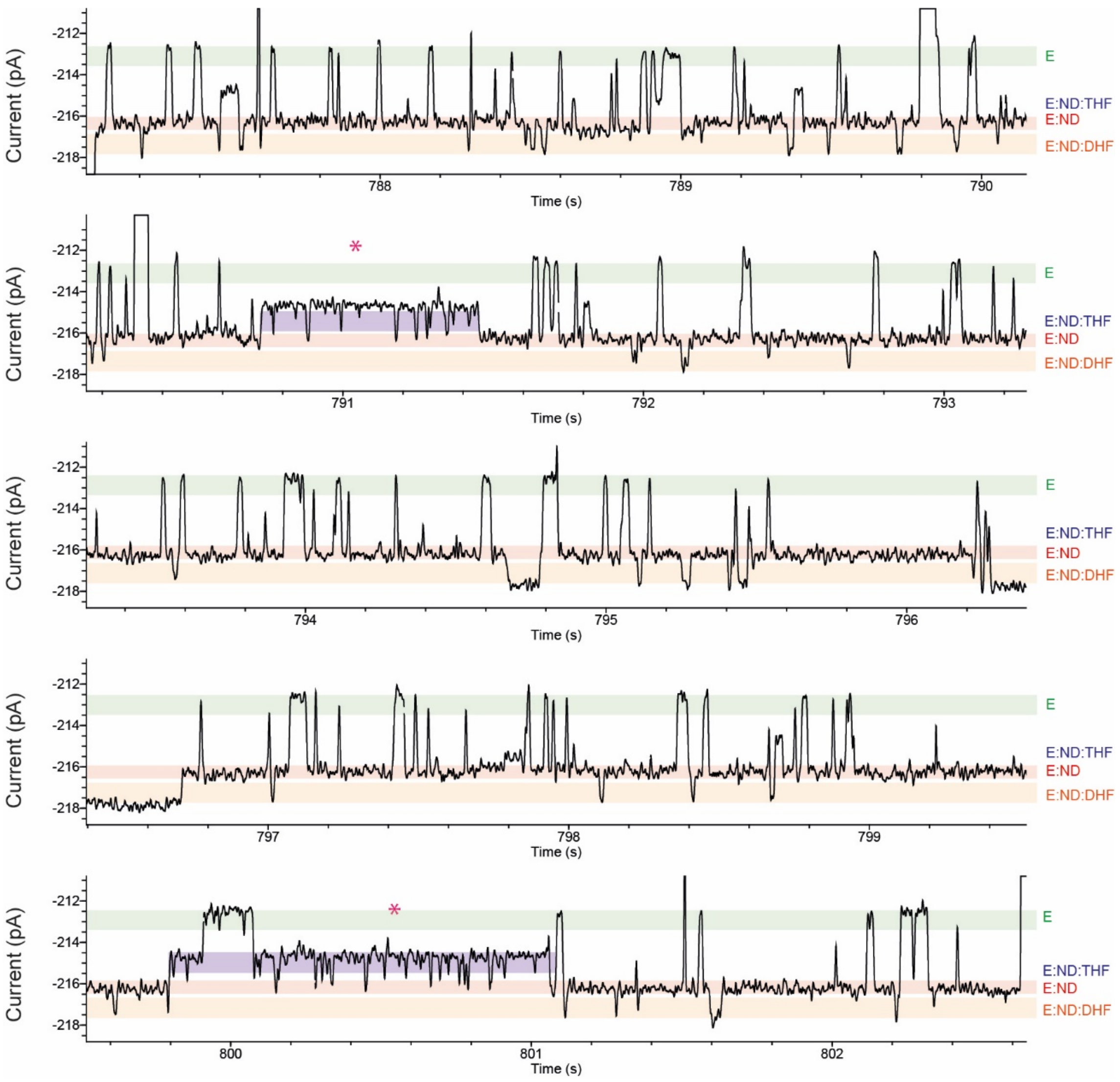


**Figure S8: Continuous recording of DHFR$_{tag}$ conformational changes during the catalysed reactions with deuterated cofactor at pH 9.1.** NADPD (0.7 mM) and dihydrofolate (DHF) (2.1 µM) were added to the *cis* and *trans* chambers, respectively. The pink asterisks are indicating the occurrence of the product, tetrahydrofolate. The current level of the *apo*-enzyme, NADPD-tetrahydrofolate-bound level, NADPD-level and ternary complex level are shown as green, purple, red and orange line, respectively. Current traces were collected in 250 mM KCl, 15 mM Tris-HCl pH 9.1 at room temperature (25 ºC), by applying a Bessel-low pass filter with a 2 kHz cut-off and sampled at 10 kHz. The trace was filtered digitally with a Gaussian low-pass filter with a 100 Hz cut-off.

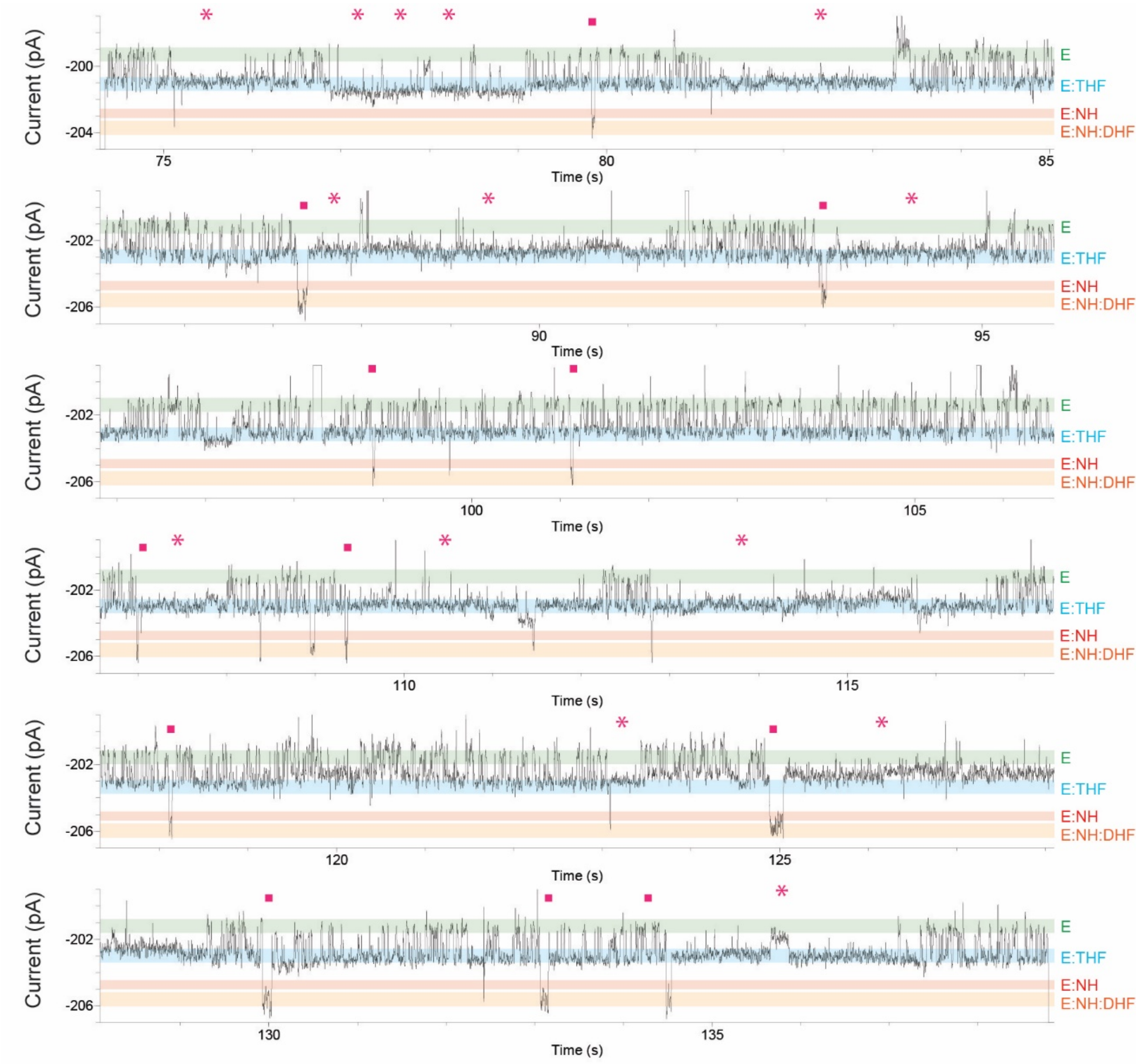


**Figure S9: Continuous recording of DHFR$_{tag}$ conformational changes during the catalysed reactions.** NADPH (5.5 μM) and dihydrofolate (DHF) (50.5 μM) were added to the *cis* and *trans* chambers, respectively. The pink squares are indicating the conformational rearrangements. The pink asterisks are indicating the occurrence of the product, tetrahydrofolate. The current level of the *apo*-enzyme, tetrahydrofolate-bound level, NADPH-level and ternary complex level are shown as green, blue, red and orange line, respectively.
